# Beyond generalization: Enhancing accurate interpretation of flexible models

**DOI:** 10.1101/808261

**Authors:** Mikhail Genkin, Tatiana A. Engel

## Abstract

Machine learning optimizes flexible models to predict data. In scientific applications, there is a rising interest in interpreting these flexible models to derive hypotheses from data. However, it is unknown whether good data prediction guarantees accurate interpretation of flexible models. We test this connection using a flexible, yet intrinsically interpretable framework for modeling neural dynamics. We find that many models discovered during optimization predict data equally well, yet they fail to match the correct hypothesis. We develop an alternative approach that identifies models with correct interpretation by comparing model features across data samples to separate true features from noise. Our results reveal that good predictions cannot substitute for accurate interpretation of flexible models and offer a principled approach to identify models with correct interpretation.

---

With advances in measurement technologies, machine learning has become critical for translating data from biological systems into theories about their function. Theories are traditionally derived by fitting data with simple *ad hoc* models, which are based on *a priori* hypotheses (*1, 2*). The best fitting model is selected and used to draw conclusions about biological mechanisms (*3–7*). An obvious pitfall is, however, that none of the *a priori* hypotheses may be correct (*8*). Alternatively, data can be fitted with flexible models, such as ar tificial neural networks (ANNs), which cover a broad class of hypotheses within a single model architecture (*9–12*) (Fig. 1). With only loose *a priori* assumptions, flexible models can discover hypotheses directly from the data. Flexible models are usually optimized for their ability to predict new data (i.e. to generalize), and the best predictive model is then analyzed and interpreted in terms of biological mechanisms (*9–14*). This widely used approach tacitly assumes that good data prediction implies correct interpretation of the model, but whether this assumption is valid is unknown. Indeed, Ptolemy’s geocentric model of the solar system predicted the movements of celestial bodies as accurately as the Copernicus’s heliocentric model. While interpreting flexible models optimized for prediction is a common practice, the derived theories can be misleading if good generalization does not guarantee correct interpretation of the model. If so, a different optimization goal is required to prioritize accurate interpretation.

**Fig 1.**
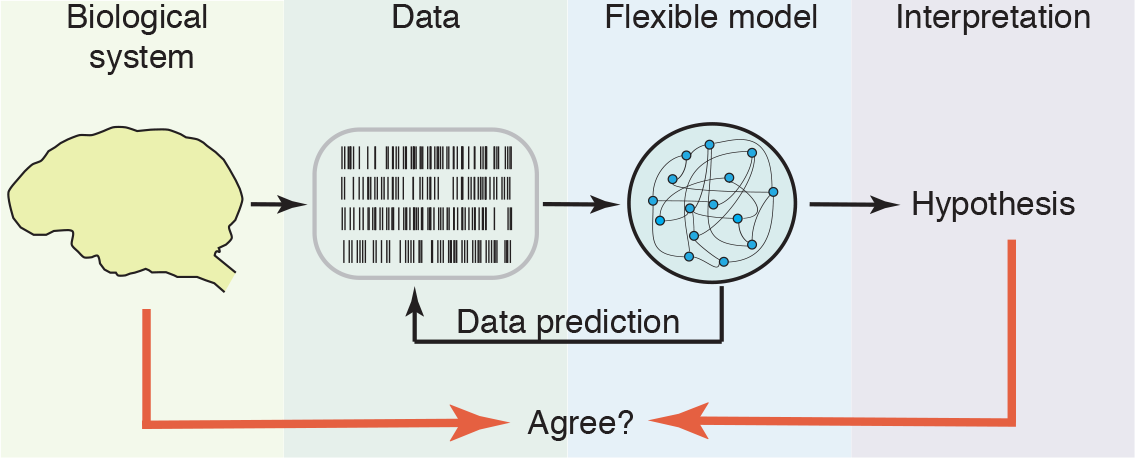
Deriving hypotheses from data using flexible models. Data is fitted with a flexible model, which covers many alternative hypotheses within a single model architecture (typically with a large number of parameters, such as in ANNs). The model is optimized for its ability to predict new data (i.e. generalize). The hypothesis is derived by interpreting the structure of the best predictive model. It is unknown whether this approach delivers correct hypotheses that accurately represent biological mechanisms.

We assessed the accuracy of interpretation of flexible models by taking advantage of a flexible, yet intrinsically interpretable framework for modeling neural dynamics (*15, 16*). This framework allows for direct comparison of the inferred hypotheses with the ground truth on synthetic data, thus testing the correctness of interpretation. As a case in point, we focus on modeling dynamics of neural responses on single trials (*3–5,8*). Inference of underlying dynamical models is notoriously hard due to doubly-stochastic nature of neural spike-trains, leading to controversial conclusions about biological mechanisms (*3–5, 8, 17*). Spikes provide sparse and irregular sampling of noisy firing-rate trajectories on single trials, which are best described as latent dynamics (*18*). Accordingly, we model spike trains as an inhomogeneous Poisson process with time-varying intensity that depends on the latent trajectory *x*(*t*) via the firing-rate function *f* (*x*) (Fig. 2A).

**Fig 2.**
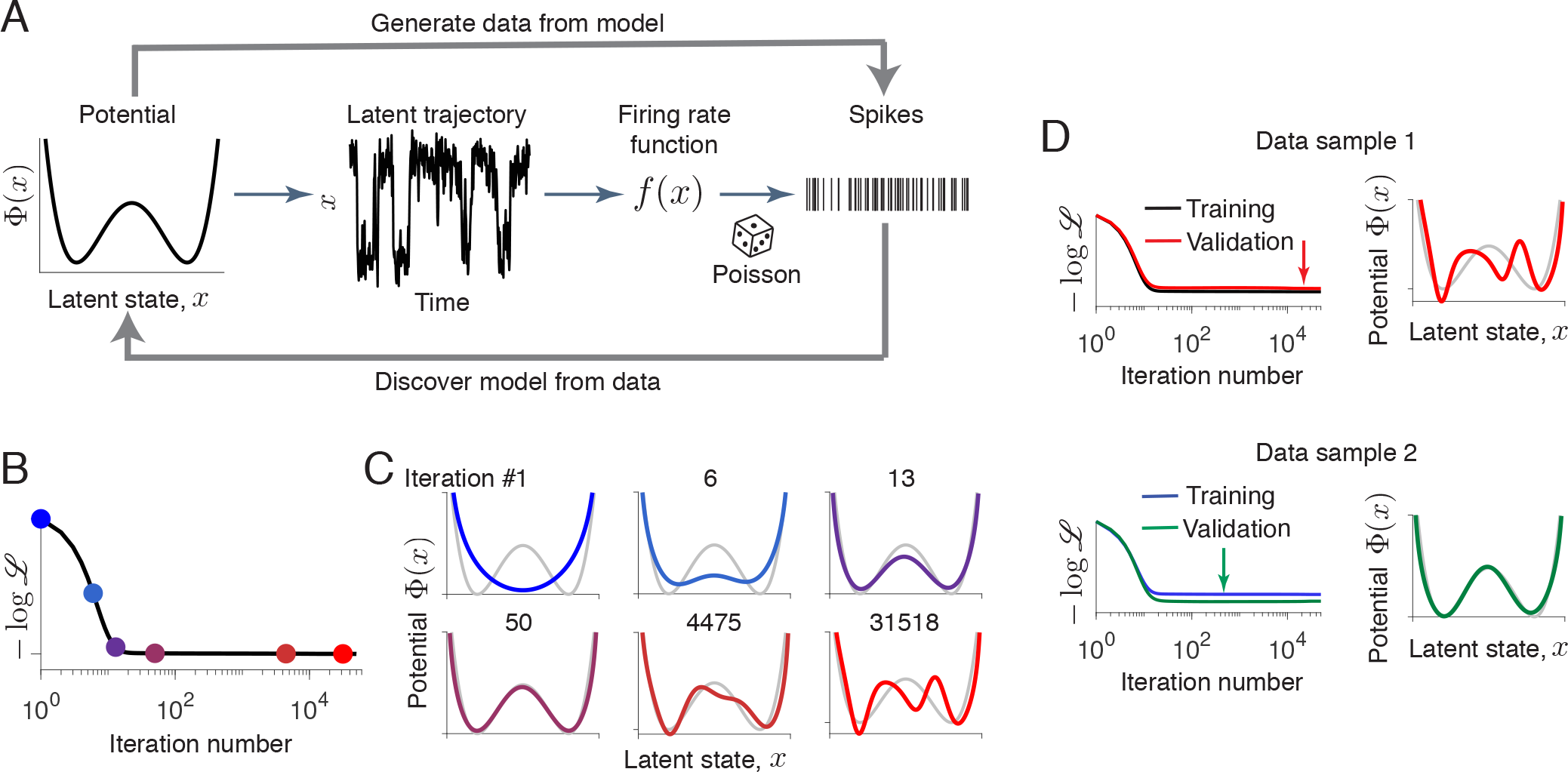
Good data prediction does not guarantee accurate interpretation of flexible models. (**A**) A flexible and intrinsically interpretable framework for modeling neural dynamics. Spike data are modeled as an inhomogeneous Poisson process with intensity that depends on the latent trajectory *x*(*t*) via firing-rate function *f* (*x*). Latent dynamics are governed by a non-linear stochastic equation (1) with a deterministic potential Φ(*x*) and Gaussian noise. (**B**) Negative log-likelihood monotonically decreases during gradient-descent optimization. (**C**) Fitted potentials Φ_*n*_(*x*) at selected iterations of the gradient-descent (colors correspond to dots in panel B) and the ground-truth potential (grey) from which the data was generated. Starting from an unspecific guess (a single-well potential on iteration 1), the optimization accurately recovers the ground-truth model (iteration 50). At later iterations, spurious features develop due to overfitting. (**D**) Left: Training and validated negative log-likelihoods for two data samples generated from the same ground-truth potential. Validated likelihood exhibits a long plateau indicating a continuum of models that generalize well. Models with the best generalization (minimum of the validated negative log-likelihood, red and green arrows) are shown to the right (colors correspond to the validated likelihood) along with the ground-truth potential (grey). The model with the best generalization matches the ground truth for data sample 2, but exhibits spurious features for data sample 1.

In our framework, the latent dynamics are governed by a non-linear stochastic equation (*15, 16*)

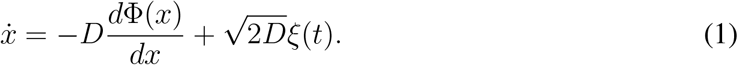

Here Φ(*x*) is a deterministic potential, and the noise *ξ*(*t*) with magnitude *D* accounts for stochasticity of latent trajectories. The potential Φ(*x*) can be any continuous function. Hence our framework covers a broad class of hypotheses, each represented by a non-linear dynamical system defined by Φ(*x*). At the same time, the shape of Φ(*x*) is intrinsically interpretable, e.g., the potential minima reveal attractors (*19*). For clarity, we focus here on inference of one-dimensional Φ(*x*) with *f* (*x*) and *D* provided (our results generalize to simultaneous inference of *F* (*x*), *f* (*x*) and *D* in multiple dimensions, materials and methods 1.7).

The hypotheses are discovered from spike data *Y* (*t*) by optimizing the shape of the potential Φ(*x*). To efficiently search through the space of all possible Φ(*x*), we developed a gradient descent optimization of the data-likelihood functional 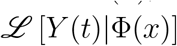 (materials and methods 1.1–1.4). We derived analytical expression for the variational derivative of log-likelihood 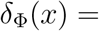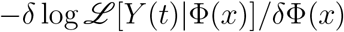. Our gradient-descent algorithm increases the data likelihood by updating the potential shape: Φ(*x*) → Φ(*x*) − γ*δ*_Φ_(*x*) (γ is a learning-rate). Similar gradient-descent algorithms are used for optimizing ANNs (*20*).

The gradient-descent optimization produces a series of dynamical models, each defined by a different potential shape Φ_*n*_(*x*) (*n* = 1, 2, … is the iteration number, Fig. 2B,C). At some intermediate iterations, the potential closely matches the ground-truth model. However, as optimization continues, Φ(*x*) develops spurious features not present in the ground-truth model. These spurious features arise mainly due to overfitting to the finite sample of stochastic spikes and are unique for each data sample. We verified that re-sampling a new data realization on each gradient-descent iteration—mimicking the infinite data regime—results in arobust recovery of the ground truth (fig. S 1). Overfitting to a finite spike sample is universally observed for different dynamics and data amount (fig. S2), which poses a challenge when ground truth is not available.

Multiple techniques exist to combat overfitting, which are all based on the idea that models matching a particular set of data too closely will fail to predict new data reliably. Accordingly, the model with the best ability to generalize is selected by evaluating its performance on a validation set of data not used for training (*18, 21*). This widely-used validation strategy aims at models that generalize well, but whether it produces models with correct interpretation (i.e. accurately matching the ground truth) is unknown. The relationship between generalization and interpretation could possibly be non-trivial for flexible models optimized on noisy data. This relationship can be effectively tested in our framework, which is difficult in ANNs that lack interpretability (i.e. the hypothesis represented by ANN’s parameters is generally unknown (*10, 22*).

We discovered that good generalization can be achieved by many models with different interpretation. We computed the validated likelihood for each model produced by the gradient-descent (Fig. 2C). The training and validated likelihoods closely track each other: after rapid initial improvement both curves level off at long plateaus. Along these plateaus, we observed a continuum of models with similar likelihood but different features. Strikingly, the plateau in the validated likelihood indicates that all models along this continuum generalize almost equally well. Indeed, spurious features develop on top of the correct potential shape that is discovered first (Fig. 2C) and have little impact on the model’s ability to predict new d ata. For example, the overfitted models generate spike trains with the first- and second-order statistics virtually indistinguishable from the ground truth (fig. S 3). Similar generalization plateaus are also observed in ANNs (fig. S4) (*23–25*).

Surprisingly, we found that selection of the model with the best generalization shows little consistency across different sets of training and validation data. The model with the best generalization is chosen at the minimum of the validated negative log-likelihood. We repeated our simulations on multiple realizations of training and validation data generated from the same ground-truth model. On some realizations, the model with the best generalization closely matches the ground truth (Fig. 2C, *lower row*). On other realizations, the model with the best generalization exhibits spurious features (Fig. 2C, *upper row*). This counterintuitive behavior arises because the validation set, just like the training set, contains noise. As a result, any model with good generalization can be chosen by chance. This problem, known as overfitting in model selection (*26*), cannot be overcome with common regularization strategies (fig. S5 and materials and methods 1.5), and it affects any flexible model optimized on finite noisy data, including ANNs (fig. S4). Although overfitting in model selection is less likely with more data, it is still substantial for realistic data amounts (Table S2). As a result, the model with the best generalization cannot be reliably interpreted.

These results entail that correct interpretation requires an optimization goal different from generalization. To identify models with correct interpretation, we leverage the fact that true features are the same, whereas noise is different across data samples. Hence, comparing models discovered on different data samples could distinguish the true features from noise. The difficulty is, however, that on different data samples, the same features are discovered at different iterations of the gradient-descent (fig. S6A). For meaningful comparisons across models, we therefore need a measure to quantify the complexity of features independent of when they are discovered. Then models of the same complexity can be directly compared for consistency of their features.

We define model complexity as a negative entropy of latent trajectories 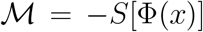 (materials and methods 1.6) (*27*). Higher model complexity indicates more structure in the potential Φ(*x*). The model complexity increases over gradient-descent iterations, as more features develop in Φ(*x*) (Fig. 3A, similar behavior is observed in ANNs (*28*)). After the true features are discovered, 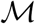 exceeds the ground-truth complexity, and further increases of 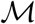 indicate fitting noise in the training data. Although the iteration when 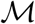 exceeds the ground-truth complexity varies across data samples (Fig. 3A), the features are aligned along the complexity axis (Fig. 3B). To detect the boundary 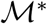 between the true features and noise, we compare models of the same complexity obtained from different data samples (Fig. 3B). For 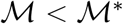, the potentials of the same complexity tightly overlap across data samples (Fig. 3C left). For 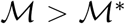, the potentials of the same complexity diverge, since overfitting patterns are unique for each data sample (Fig. 3C right). The potentials with complexity 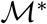 closely match the ground-truth model (Fig. 3C middle). We confirmed that complexity boundary 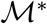 reliably indicates the model with correct interpretation for different ground-truth dynamics (fig. S6, S7). The complexity boundary 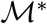 can be used to identify the model with correct interpretation in biological data when the ground truth is unknown.

**Fig 3.**
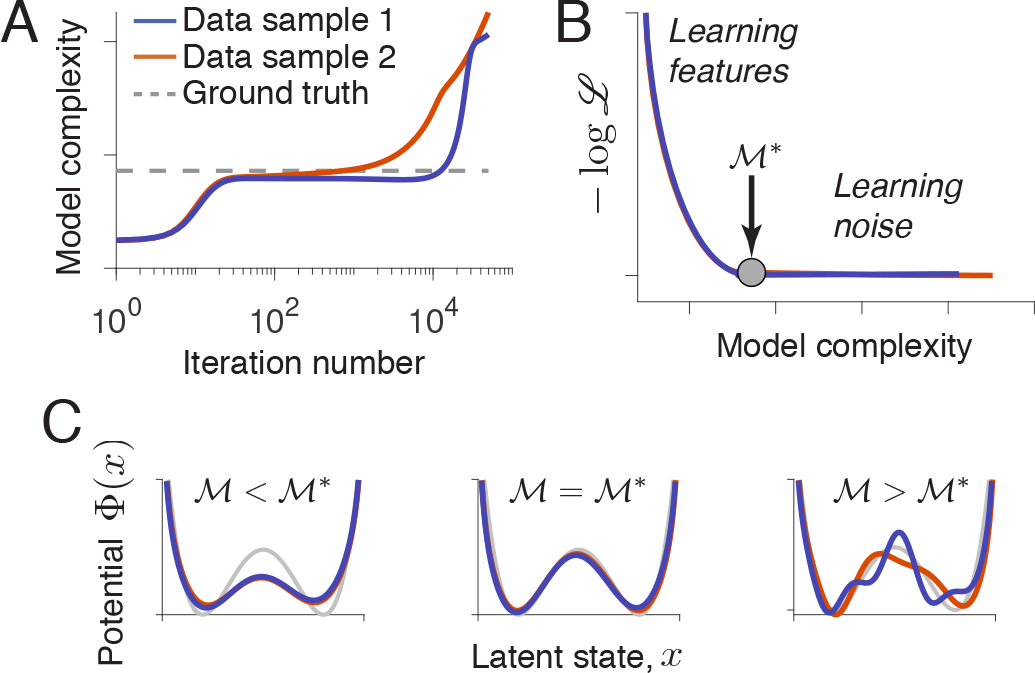
Identifying models with correct interpretation by comparing features across data samples. (**A**) Model complexity increases over gradient-descent iterations at a rate varying across data samples. The ground-truth model complexity is exceeded at different iterations for different data samples. (**B**) Normalized validated negative log-likelihood plotted against model complexity (colors correspond to data in A). 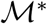 (arrow) is the maximal model complexity for which fitted potentials are consistent across data samples. 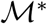 coincides with the ground-truth model complexity (dot). (**C**) Potentials for 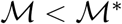 (left), 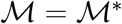 (middle) and 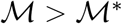 (right), colors correspond to data in A.

We validated our results using recordings of spiking activity from the visual cortex of behaving monkeys (dataset from Ref. (*2*)). In these data, neural activity spontaneously transitions between episodes of vigorous (On) and faint (Off) spiking that are irregular within and across trials (Fig. 4A). These endogenous On-Off dynamics were previously fitted with a model that assumes abrupt transitions between discrete On and Off states, and with an alternative model, which assumes smooth activity fluctuations (*2*). These *ad hoc* models with contrasting assumptions can both segment spiking activity into On and Off episodes, but they cannot resolve whether the cortical On-Off dynamics constitute fluctuations around a single attractor or transitions between multiple metastable states. Within our framework, these alternative hypotheses correspond to potential shapes with a single or multiple wells (fig. S8).

**Fig 4.**
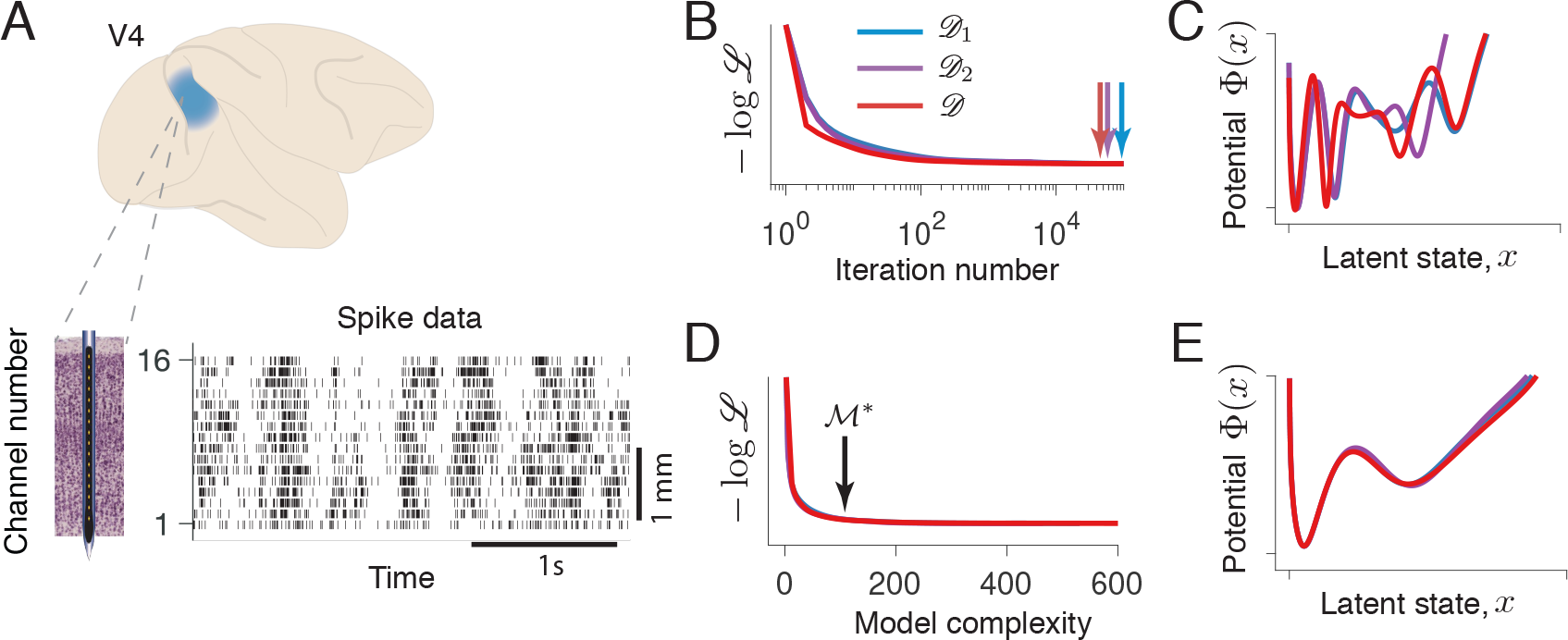
Discovering interpretable model of neural dynamics from neurophysiological recordings. (**A**) An example trial (right) showing spontaneous transitions between episodes of vigorous (On) and faint (Off) spiking in multiunit activity simultaneously recorded with 16-channel electrodes (left) from the primate visual cortical area V4 during a fixation task. Spikes are marked by vertical ticks. Modeling results for the first channel are shown in B-E. (**B**) Validated negative log-likelihoods over iterations of the gradient-descent for three data samples 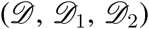. Models with the best generalization correspond to minima indicated by arrows. (**C**) Models with the best generalization are inconsistent across data samples. (**D**) Validated negative log-likelihoods plotted against model complexity for three data samples 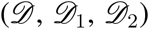. 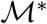 (arrow) is the maximal model complexity for which potentials are consistent across data samples. (**E**) The potentials at 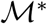 tightly overlap for three data samples. The potential shape supports the hypothesis of metastable transitions. Colors in panels C-E correspond to data in B.

We divide the full data 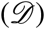 in halves (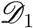 and 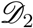), and further divide each set 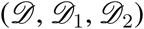 in the training and validation sets. We perform gradient-descent optimization on each training set, evaluate models on the corresponding validation set, and track consistency of features in models discovered from 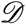, 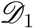, and 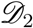 along the complexity axis to find 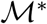. We fit spikes on each recorded channel separately (example channel in Fig. 4B-E), and compare model fits across all channels (fig. S9). Since On-Off dynamics are largely synchronous across the population (*2*), similarity of potentials discovered from different channels would indicate that our approach reliably identifies models with correct interpretation.

With the neurophysiological recordings, we observe the same phenomena as described for synthetic data. The gradient-descent optimization continuously improves the training likelihood, producing a sequence of models with increasing complexity. Many of these models generalize equally well, which manifests in a long plateau in the validated likelihood (Fig. 4B). The models with the best generalization are inconsistent across datasets 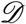, 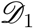, and 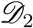 (Fig. 4C), indicating likely overfitting in model selection. The complexity boundary *∗* reliably identifies models that are consistent across datasets 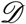, 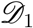, and 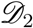 (Fig. 4D) and across channels (fig. S9). The inferred potentials exhibit two wells, suggesting that On-Off dynamics are metastable transitions and not fluctuations around a single attractor.

We leverage a flexible and intrinsically interpretable framework to demonstrate that good data prediction does not guarantee correct interpretation of flexible models. Gradient-descent optimization discovers many models that generalize well despite differences in their features and interpretation. Overfitting in model selection affects any flexible model fitted and validated on finite noisy data. Our results raise a caution for methods based on fitting data with a flexible model (e.g., ANN) and interpreting the fitted model as a biological mechanism. Flexible models require non-trivial hyperparameter tuning geared towards models with the best generalization, and our results suggest that interpretation of such models may be uncertain. Our work is the first to explore the link b etween generalization a nd i nterpretation, which have been studied only separately in ANNs that lack interpretability. Models with correct interpretation can be reliably identified by comparing features of the same complexity discovered on different data samples. This comparison requires quantifying the complexity of fitted models, in contrast to conventional measures of complexity for ANNs that characterize the full capacity of the model architecture. Developing appropriate complexity measures for ANNs is a significant outstanding issue, the solution of which would provide the necessary theoretical foundation for interpretable machine learning.

## Acknowledgements

This work was supported by the NIH grant R01 EB026949 and the Swartz Foundation. We thank G. Angeris for help at early project stages, K. Haas for useful discussions, and P. Koo, J. Jansen, A. Siepel, J. Kinney, and T. Janowitz for their thoughtful comments on the manuscript. We thank N.A. Steinmetz and T. Moore for sharing the electrophysiological data, which are presented in (*2*) and are archived at the Stanford Neuroscience Institute server at Stanford University.

## Author Contributions

M.G. and T.A.E. designed the study, performed research, discussed the findings and wrote the paper.

## Supplementary Materials

Materials and Methods

Figures S1–S9

Tables S1–S2

References (*29–32*)

