## Supplementary Information for "Beyond generalization: Enhancing accurate interpretation of flexible models"

### Supplementary materials for: Beyond generalization: Enhancing accurate interpretation of flexible models

October 16, 2019

**3 Supplementary figures** **15**

### 1 Materials and methods

#### 1.1 Maximum likelihood inference

In the non-parametric framework, the spiking activity depends on the latent trajectory $x(t)$ via the firing-rate function $f(x)$. The latent dynamics are controlled by the deterministic potential $\Phi(x)$ and noise with the magnitude $D$ (Fig. 2 in the main text). Here we focus on inference of

$\Phi(x)$ from the data, assuming that $D$ and $f(x)$ are known. We infer the potential $\Phi(x)$ from data $Y(t)$ by maximizing the data likelihood:

$$\tilde{\Phi}(x) = \underset{\Phi(x)}{\operatorname{argmax}}\, \mathscr{L}\left[Y(t)|\Phi(x)\right]. \tag{1}$$

The likelihood functional is a probability that the data $Y(t)$ comes from the given model $\Phi(x)$:

$$\mathscr{L}\left[Y(t)|\Phi(x)\right] = P(Y(t)|\Phi(x)). \tag{2}$$

The optimization problem in Eq. (1) is solved iteratively using gradient-descent (GD) algorithm. On each GD iteration, we need to calculate both the likelihood $\mathscr{L}\left[Y(t)|\Phi(x)\right]$ and its variational derivative $\delta\mathscr{L}\left[Y(t)|\Phi(x)\right]/\delta\Phi(x)$. We derive analytical expressions for the likelihood and its variational derivative, which we evaluate numerically on each GD iteration. Rather than updating the potential $\Phi(x)$, we update the driving force $F(x) = -d\Phi(x)/dx$, for which we calculate the variational derivative $\delta\mathscr{L}\left[Y(t)|\Phi(x)\right]/\delta F(x)$. The potential $\Phi(x)$ is calculated from the force $F(x)$ by taking an antiderivative.

#### 1.2 Likelihood calculation

Analytical formula for the likelihood calculation is briefly outlined here (see [15] for additional details). The likelihood (2) is calculated via marginalization of the joint probability density $P(X(t), Y(t)|\Phi(x))$ of the observed spike-data $Y(t)$ and the latent trajectory $X(t)$:

$$\mathscr{L}\left[Y(t)|\Phi(x)\right] = \int \mathscr{D}X(t)\; P(X(t), Y(t)|\Phi(x)). \tag{3}$$

The brute-force evaluation of Eq. (3) would require intractable integration over all possible latent paths. This calculation can be simplified using the Markov property of the latent dynamics and conditional independence of observations. For the Langevin dynamics (Eq. (1) in the main text), the transition probability density over the latent space $p(x_{t_k}, t_k|x_{t_{k-1}}, t_{k-1})$ does not depend on the intermediate states, and spikes are independent when conditioned on the latent states. The joint probability density $P(X(t), Y(t))$ can therefore be factorized into a product of probability densities of spike observations and transition probability densities over the latent space between the adjacent spikes:

$$P(X(t), Y(t)) = p(x_{t_0}) \prod_{k=1}^{N} p(y_{t_i}|x_{t_i}) p(x_{t_i}|x_{t_{i-1}}). \tag{4}$$

Here $t_1, t_2, ..., t_N$ are the observed spike times and $t_0$ is the experiment onset time. $X(t) = \{x_{t_0}, x_{t_1}, ..., x_{t_N}\}$ is a latent trajectory at these times. $Y(t) = \{y_{t_1}, y_{t_2}, ..., y_{t_N}\}$ are the observed spikes. $p(y_{t_i}|x_{t_i})$ is the probability of observing a spike at time $t_i$ given the latent position $x_{t_i}$. Finally, $p(x_{t_i}|x_{t_{i-1}})$ is the transition probability density over the latent space from $x_{t_{i-1}}$ to $x_{t_i}$ during the time between adjacent spike observations. Note that $X(t)$ is a discretized latent path sampled at $N+1$ time points, and all other time points are absorbed within the transition probability densities. The likelihood is calculated from Eq. (4) via marginalization over the latent space variables:

$$\mathscr{L} = \int_{x_{t_0}} \int_{x_{t_1}} \cdots \int_{x_{t_N}} dx_{t_0} dx_{t_1} \ldots dx_{t_N} P(X(t), Y(t)). \tag{5}$$

The transition probability density over the latent space is a solution of the generalized Fokker-Planck equation [15]:

$$\frac{\partial p(x,t)}{\partial t} = \left(-D\frac{\partial}{\partial x}F(x) + D\frac{\partial}{\partial x^2} - f(x)\right) p(x,t) \equiv -\hat{\boldsymbol{\mathcal{H}}} p(x,t). \tag{6}$$

The first two terms in the operator $\hat{\mathcal{H}}$ are the usual drift and diffusion terms that originate from the Langevin dynamics (Eq. (1) in the main text). The last term $-f(x)$ accounts for the fact that no spikes were observed between each of the time points $t_k$ and $t_{k-1}$.

Following [15], we transform Eq. (6) in a Hermitian form with the operator $\mathcal{H}$. This operator is defined as $\mathcal{H} = \exp(\Phi(x)/2)\hat{\mathcal{H}}\exp(-\Phi(x)/2)$. Introducing a new variable $\rho(x,t) = p(x,t)\exp(\Phi(x)/2)$, one can show that Eq. (6) is transformed into:

$$\frac{\partial \rho(x,t)}{\partial t} = -\mathcal{H}\rho(x,t). \tag{7}$$

We will use Eq. (7) to evaluate $\rho(x,t)$ over time. The original probability density $p(x,t)$ can be recovered by $p(x,t) = \rho(x,t)\exp(-\Phi(x)/2)$. We also introduce the equilibrium probability density $p_{\text{eq}}(x) = \exp(-\Phi(x))$, where we choose the arbitrary scaling of the potential $\Phi(x)$ so that $p_{\text{eq}}(x)$ is normalized. Then $p(x,t) = \rho(x,t)\sqrt{p_{\text{eq}}(x)}$.

The formal solution of Eq. (7) can be written as the operator exponential:

$$\rho(x_{t_i}|x_{t_{i-1}}) = e^{-\mathcal{H}\Delta t_i}, \tag{8}$$

where $\Delta t_i = t_i - t_{i-1}$. To compute the operator $e^{-\mathcal{H}\Delta t_i}$ efficiently for any $\Delta t_i$, we use the eigenfunctions $\Psi(x)$ and eigenvalues $\lambda$ of the operator $\mathcal{H}$, which satisfy

$$\mathcal{H}\Psi(x) = \lambda\Psi(x). \tag{9}$$

We solve this eigenvalue problem numerically. To this end, we discretize the continuous domain $x$ using the Spectral Elements Method (SEM), and solve for the eigenvalues $\lambda_i$ and the eigenvectors $Q_{ij} = \Psi_j(x_i)$ of the discretized operator $\mathcal{H}$ (see Section 2 for details). The matrix $\boldsymbol{Q}$ contains all eigenvectors of the discretized $\mathcal{H}$. As in [15], we truncate the basis $\boldsymbol{Q}$ to the 64 eigenvectors corresponding to the 64 largest eigenvalues. To confirm that this number is sufficient, we also used 128 eigenvectors and observed identical results.

In the eigenbasis $\boldsymbol{Q}$ of the operator $\mathcal{H}$, any continuous function $g(x)$ is represented by a vector $\boldsymbol{g}$ with elements equal to the expansion coefficients in this basis:

$$g_i = \int_x g(x)\Psi_i(x)dx. \tag{10}$$

The operator exponential $e^{-\mathcal{H}\Delta t_k}$ is represented by a transition matrix $\boldsymbol{A}_k$ with the elements:

$$A_{k,ij} = \int_x \Psi_i(x)e^{-\mathcal{H}\boldsymbol{\Delta t_k}}\Psi_j(x)dx = \delta_{ij}e^{-\lambda_i\Delta t_k}, \tag{11}$$

where we took advantage of the eigenbasis properties.

The probability density of a spike emission at a given latent state $p(y_{t_i}|x_{t_i})$ is given by $p(y|x) = f(x)$, just by the definition of the firing rate. The corresponding emission operator is represented by the matrix $\boldsymbol{B}$ in the basis $\boldsymbol{Q}$, which is calculated using the rules of basis transformations:

$$\boldsymbol{B} = \boldsymbol{Q^T W f Q}. \tag{12}$$

Here the diagonal matrix $\boldsymbol{f}$ represents the emission operator in the SEM basis, and $\boldsymbol{W}$ is a diagonal matrix of SEM weights (see Section 2).

With the developed formalism, Eq. (5) turns into a chain of matrix multiplications, where matrix-matrix products automatically account for the marginalization over the intermediate variables $x_{t_0}, x_{t_2}, ...x_{t_{N-1}}$. To marginalize over the last latent position $x_{t_N}$, we multiply this chain by a column vector $\boldsymbol{\beta_N}$:

$$\mathscr{L} = \boldsymbol{\alpha}_0^T\boldsymbol{A}_1\boldsymbol{B}\boldsymbol{A}_2\boldsymbol{B}...\boldsymbol{A}_N\boldsymbol{B}\boldsymbol{\beta_N}. \tag{13}$$

Here $\boldsymbol{\alpha}_0 = \boldsymbol{Q}^T \boldsymbol{W} \boldsymbol{\rho_0}$ is a vector representing the initial probability distribution transformed into the basis $\boldsymbol{Q}$ (the vector $\boldsymbol{\rho_0}$ is a discretization of $\rho(x, 0)$ in the SEM basis). The initial condition is chosen to be the equilibrium probability distribution, $p(x, 0) = p_{\text{eq}}(x)$, so that $\rho(x, 0) = \sqrt{p_{\text{eq}}(x)} = \rho_{\text{eq}}(x)$. The terminal vector $\boldsymbol{\beta_N}$ is defined by the probability normalization condition (similar to the Hidden Markov Model, HMM [18]). Since

$$\frac{p(Y(t), x_N)}{\sqrt{p_{\text{eq}}(x)}} = \boldsymbol{\alpha}_0^T \boldsymbol{A}_1 \boldsymbol{B} \boldsymbol{A}_2 \boldsymbol{B} ... \boldsymbol{A}_N \boldsymbol{B}, \tag{14}$$

we see that in order to satisfy the condition $\mathscr{L} = \int_{x_N} p(Y(t), x_N) dx$, we need to set $\boldsymbol{\beta_N} = \boldsymbol{Q}^T \boldsymbol{W} \boldsymbol{\rho_{\text{eq}}}$, which follows from the normalization conditions for the probability density $p_{\text{eq}}(x)$.

In summary, given the model specified by $\theta = \{\Phi(x), f(x), D\}$, the likelihood is calculated using Eq. (13), where the operator matrices are found from Eqs. (11),(12) with the eigenvectors and eigenvalues of operator $\boldsymbol{\mathcal{H}}$ obtained from Eq. (9). More details on numerical implementation are given in Section 2.

In addition to matrix-vector form, for convenience we also use Dirac bra-ket notation. In this notation, left-vectors correspond to bra-states $\langle\rho|$, right vectors to ket-state $|\rho\rangle$, and matrices to operators. With a choice of basis, the states are transformed into vectors and operators into matrices, so that both formalisms are equivalent. In Dirac notation Eq. (13) can be written as a chain of operators that propagate initial state $\langle\alpha_0|$ towards the terminal state $|\beta_N\rangle$:

$$\mathscr{L}\left[Y(t), \Phi(x)\right] = \left\langle \alpha_0 \middle| e^{-\boldsymbol{\mathcal{H}}\Delta t_1} \boldsymbol{y}_{t_1} e^{-\boldsymbol{\mathcal{H}}\Delta t_2} \boldsymbol{y}_{t_2} ... e^{-\boldsymbol{\mathcal{H}}\Delta t_N} \boldsymbol{y}_{t_N} \middle| \beta_N \right\rangle, \tag{15}$$

where $\boldsymbol{y}_{t_k}$ is the spike emission operator.

We evaluate the likelihood using the forward-backward algorithm similar to that for HMMs [18]. Starting from the initial state $\alpha_0$, the states $\alpha_1, \alpha_2, ..., \alpha_N$ are calculated recursively through the forward pass:

$$\langle\alpha_n| = \langle\alpha_{n-1}| e^{-\boldsymbol{\mathcal{H}}\Delta t_n} \boldsymbol{y}_{t_n}, \qquad n = 1, 2, ..., N. \tag{16}$$

From Eq. (15), the likelihood $\mathscr{L} = \langle\alpha_N|\beta_N\rangle$. The subsequent backward pass is needed for evaluation the likelihood gradients:

$$|\beta_{n-1}\rangle = e^{-\boldsymbol{\mathcal{H}}\Delta t_n} \boldsymbol{y}_{t_n} |\beta_n\rangle, \qquad n = 1, 2, ..., N. \tag{17}$$

#### 1.3 Variational derivatives of the likelihood functional

We derive the gradient-descent algorithm for maximizing the likelihood. In contrast to the previous work [15], which used an approximate expectation-maximization algorithm, the gradient-descent update rule can be derived exactly. This update requires to compute the variational derivative $\delta\mathscr{L}\left[Y(t)|\Phi(x)\right]/\delta\Phi(x)$. For numerical stability, rather than updating the potential $\Phi(x)$, we update the driving force $F(x)$ using $\delta\mathscr{L}/\delta F(x)$. The dependence of likelihood on $F(x)$ is hidden in the operator $\boldsymbol{\mathcal{H}}$, see Eq. (15). Using the product rule, the variational derivative can be written as:

$$\frac{\delta\mathscr{L}}{\delta F(x)} = \sum_{\tau=1}^{N} \sum_{i,j} a_i(\tau) b_j(\tau+1) \frac{\delta\langle\Psi_i| e^{-\boldsymbol{\mathcal{H}}\Delta t_\tau} |\Psi_j\rangle}{\delta F} + \frac{\delta\langle\alpha_0|}{\delta F}|\beta_0\rangle + \langle\alpha_N|\frac{\delta|\beta_N\rangle}{\delta F}. \tag{18}$$

Here $\Psi_i$ are the eigenvectors of $\boldsymbol{\mathcal{H}}$, and we introduced the notation:

$$a_i(\tau) = \langle\alpha_\tau|\Psi_i\rangle, \qquad b_j\tau) = \langle\Psi_j|\beta_\tau\rangle. \tag{19}$$

For numerical stability, it is more convenient to optimize the log-likelihood $\log \mathscr{L}$, which has the derivative

$$\frac{\delta \log \mathscr{L}}{\delta F(x)} = \frac{1}{\mathscr{L}} \frac{\delta \mathscr{L}}{\delta F(x)}. \tag{20}$$

In the following discussion we omit the factor $1/\mathscr{L}$ for convenience and derive the expression for $\delta \mathscr{L}/\delta F(x)$.

The first term on the right-hand side of Eq. (18) can be calculated using the formula for derivative of an exponentiated operator [29]:

$$\frac{\partial}{\partial \lambda} e^{-\beta \boldsymbol{H}} = -\int_0^\beta e^{-(\beta-u)\boldsymbol{H}} \frac{\delta \boldsymbol{H}}{\delta \lambda} e^{-u\boldsymbol{H}} du. \tag{21}$$

Using Eq. (21), one can derive:

$$\begin{aligned} \frac{\delta \langle \Psi_i | e^{-\boldsymbol{\mathcal{H}}\Delta t_\tau} | \Psi_j \rangle}{\delta F} &= -\int_0^{\Delta t_\tau} \left\langle \Psi_i \middle| e^{-(\Delta t_\tau - u)\boldsymbol{\mathcal{H}}} \frac{\delta \boldsymbol{\mathcal{H}}}{\delta F} e^{-u\boldsymbol{\mathcal{H}}} \middle| \Psi_j \right\rangle du = \\ = -\sum_{l,m} \int_0^{\Delta t_\tau} &\left\langle \Psi_i \middle| e^{-(\Delta t_\tau - u)\boldsymbol{\mathcal{H}}} \middle| \Psi_l \right\rangle \left\langle \Psi_l \middle| \frac{\delta \boldsymbol{\mathcal{H}}}{\delta F} \middle| \Psi_m \right\rangle \left\langle \Psi_m \middle| e^{-u\boldsymbol{\mathcal{H}}} \middle| \Psi_j \right\rangle du = \\ = -\sum_{l,m} \int_0^{\Delta t_\tau} &\langle \Psi_i | \Psi_l \rangle e^{-(\Delta t_\tau - u)\lambda_i} \left\langle \Psi_l \middle| \frac{\delta \boldsymbol{\mathcal{H}}}{\delta F} \middle| \Psi_m \right\rangle \langle \Psi_m | \Psi_j \rangle e^{-u\lambda_j} du = \\ = -\left\langle \Psi_i \middle| \frac{\delta \boldsymbol{\mathcal{H}}}{\delta F} \middle| \Psi_j \right\rangle &\int_0^{\Delta t_\tau} e^{-(\Delta t_\tau - u)\lambda_i} e^{-u\lambda_j} du \equiv -\left\langle \Psi_i \middle| \frac{\partial \boldsymbol{\mathcal{H}}}{\partial F} \middle| \Psi_j \right\rangle \Gamma^\tau_{i,j}. \end{aligned} \tag{22}$$

Here $\lambda$ are the eigenvalues of $\boldsymbol{\mathcal{H}}$, and $\Gamma^\tau_{i,j}$ is defined as [15]:

$$\Gamma^\tau_{ij} = \int_0^{\Delta t_\tau} e^{-(\Delta t_\tau - u)\lambda_i} e^{-u\lambda_j} du = \begin{cases} \Delta t_\tau e^{-\lambda_i \Delta t_\tau}, & i = j; \\ \frac{e^{-\lambda_i \Delta t_\tau} - e^{-\lambda_j \Delta t_\tau}}{\lambda_j - \lambda_i}, & i \neq j. \end{cases} \tag{23}$$

The term $\left\langle \Psi_i \middle| \frac{\partial \boldsymbol{\mathcal{H}}}{\partial F} \middle| \Psi_j \right\rangle$ can be calculated using the Euler-Lagrange equation. The operator $\boldsymbol{\mathcal{H}}$ is written in terms of $F(x)$ as:

$$\boldsymbol{\mathcal{H}} = -D\nabla^2 + D\frac{F'(x)}{2} + D\frac{F^2(x)}{4} - f(x). \tag{24}$$

Accordingly,

$$\left\langle \Psi_i \middle| \frac{\delta \boldsymbol{\mathcal{H}}}{\delta F} \middle| \Psi_j \right\rangle = \frac{\delta}{\delta F} D \int \Psi_i(x) \left( \frac{F'(x)}{2} + \frac{F(x)^2}{4} \right) \Psi_j(x) dx = \frac{D}{2} \left( F(x)\Psi_i(x)\Psi_j(x) - \frac{d\left(\Psi_i(x)\Psi_j(x)\right)}{dx} \right). \tag{25}$$

This expression can be simplified:

$$\left\langle \Psi_i \middle| \frac{\delta \boldsymbol{\mathcal{H}}}{\delta F} \middle| \Psi_j \right\rangle = -\frac{D}{2} p_{eq}(x) \frac{d\left(\phi_i(x)\phi_j(x)\right)}{dx}, \tag{26}$$

where $\phi(x) = \Psi(x)/\sqrt{p_{eq}(x)}$.

The last two terms on the r.h.s. of Eq. (18) can be calculated from the definitions of $\langle \alpha_0 |$ and $| \beta_N \rangle$ using the chain rule for functional derivatives. In the basis of $\boldsymbol{\mathcal{H}}$ we have $\langle \alpha_0 | = | \beta_N \rangle = \sqrt{p_{eq}(x)} = \rho_{eq}(x)$. Introducing:

$$\mathscr{L}_{AB} = \int_{-1}^{1} dx (\beta_0(x) + \alpha_N(x)) \rho_{\mathrm{eq}}(x) dx, \tag{27}$$

we need to find $\delta\mathscr{L}_{AB}/\delta F(x)$, where the dependence of $\beta_0(x)$ and $\alpha_N(x)$ on $F(x)$ is ignored, since it was already taken into account in other terms in Eq. (18). Using the chain rule [30], we obtain:

$$\frac{\delta\mathscr{L}_{\rm AB}}{\delta F(y)} = \int_{-1}^{1} ds \frac{\delta\mathscr{L}_{\rm AB}}{\delta p_{eq}[s]} \frac{\delta p_{eq}[s]}{\delta F(y)}. \tag{28}$$

Using the Euler-Lagrange equation, we obtain for the first term:

$$\frac{\delta\mathscr{L}_{\rm AB}}{\delta p_{eq}[s]} = \frac{\beta_0(s)+\alpha_N(s)}{2\rho_{\rm eq}(s)}. \tag{29}$$

To calculate the second term, we need to express $p_{eq}$ in terms of the force:

$$p_{\rm eq} = \frac{\exp(\int_{-1}^{x} F(x')dx')}{\int_{-1}^{1}\exp(\int_{-1}^{x} F(x')dx')dx} \equiv \frac{e^{G(x)}}{\int_{-1}^{1} e^{G(x')}dx'} = \frac{\int_{-1}^{1} e^{G(x')}\delta(x-x')dx'}{\int_{-1}^{1} e^{G(x')}dx'}. \tag{30}$$

Here $G(x) = \int_{-1}^{1} F(x')H(x-x')dx'$, and $H$ is a Heaviside function. Using the chain rule again, we obtain:

$$\frac{\delta p_{eq}[s]}{\delta F(y)} = \int_{-1}^{1} ds' \frac{\delta p_{eq}[s]}{\delta G[s']}\frac{\delta G[s']}{\delta F(y)}. \tag{31}$$

Equating these expressions with the Euler-Lagrange rule, we get:

$$\begin{aligned} \frac{\delta p_{eq}[s]}{\delta G[s']} &= \frac{e^{G(s')}\delta(s-s')\int_{-1}^{1} e^{G(x')}dx' - e^{G(s')}e^{G(s)}}{\left(\int_{-1}^{1} e^{G(x')}dx'\right)^2}, \\ \frac{\delta G[s']}{\delta F(y)} &= H(s'-y). \end{aligned} \tag{32}$$

In result,

$$\begin{aligned} \frac{\delta p_{eq}[s]}{\delta F(y)} &= \int_{-1}^{1} ds' \left[\frac{e^{G(s')}\delta(s-s')}{\int_{-1}^{1} e^{G(x')}dx'} - \frac{e^{G(s')}e^{G(s)}}{\left(\int_{-1}^{1} e^{G(x')}dx'\right)^2}\right] H(s'-y) = \\ &= \frac{e^{G(s)}}{\int_{-1}^{1} e^{G(x')}dx'}\left[H(s-y) - \frac{\int_{-1}^{1} e^{G(s')}H(s'-y)ds'}{\int_{-1}^{1} e^{G(x')}dx'}\right] = p_{eq}(s)\left[H(s-y) - \frac{\int_{y}^{1} e^{G(s')ds'}}{\int_{-1}^{1} e^{G(x')}dx'}\right] = \\ &= p_{eq}(s)\left[H(s-y) - \int_{y}^{1} p_{eq}(s')ds'\right] = p_{eq}(s)\left[H(s-y) - 1 + \int_{-1}^{y} p_{eq}(s')ds'\right]. \end{aligned} \tag{33}$$

Substituting Eqs. (29),(33) into Eq. (28), we obtain:

$$\begin{aligned} \frac{\delta\mathscr{L}_{\rm AB}}{\delta F(y)} &= \int_{-1}^{1} ds \frac{\beta_0(s)+\alpha_N(s)}{2\rho_{\rm eq}(s)} p_{eq}(s)\left[H(s-y) - 1 + \int_{-1}^{y} p_{eq}(s')ds'\right] = \\ &= \frac{1}{2}\int_{-1}^{1} ds\,(\beta_0(s)+\alpha_N(s))\,\rho_{\rm eq}(s)\left[H(s-y) - 1 + \int_{-1}^{y} p_{eq}(s')ds'\right] = \\ &= \frac{1}{2}\left[\int_{y}^{1}(\beta_0(s)+\alpha_N(s))\,\rho_{\rm eq}(s)ds + \left(\int_{-1}^{1} ds\,(\beta_0(s)+\alpha_N(s))\,\rho_{\rm eq}(s)\right)\left(-1 + \int_{-1}^{y} p_{eq}(s')ds'\right)\right] = \\ &= \frac{1}{2}\left[2 - \int_{-1}^{y}(\beta_0(s)+\alpha_N(s))\,\rho_{\rm eq}(s)ds - 2 + 2\int_{-1}^{y} p_{eq}(s)ds\right] = \int_{-1}^{y}\left(p_{eq}(s) - \frac{\beta_0(s)+\alpha_N(s)}{2}\rho_{\rm eq}(s)\right)ds. \end{aligned} \tag{34}$$

Here we used the property

$$\int_{-1}^{1} ds\beta_0(s)\rho_{\text{eq}}(s) = \int_{-1}^{1} ds\alpha_N(s)\rho_{\text{eq}}(s) = 1, \tag{35}$$

which follows from the probability normalization condition and scaling of $\alpha$ and $\beta$ coefficients.

The final expression for the variational derivative reads:

$$\frac{\delta\mathscr{L}}{\delta F} = \sum_{ij} G_{ij}\frac{D}{2}p_{eq}(x)\frac{d\left(\phi_i(x)\phi_j(x)\right)}{dx} + \int_{-1}^{y}\left(p_{eq}(s) - \frac{\beta_0(s)+\alpha_N(s)}{2}\rho_{\text{eq}}(s)\right)ds, \tag{36}$$

where we introduced:

$$G_{ij} = \sum_{\tau}\Gamma_{i,j}^{\tau}\alpha_i(\tau)\beta_j(\tau+1). \tag{37}$$

First, the function $G_{ij}$ is calculated during the backward pass Eq. (17), and after that Eq. (36) is evaluated. Calculation of an antiderivative in the SEM basis is described in Section 2.

#### 1.4 Gradient-descent optimization

In the gradient-descent optimization, we start with an initial guess of the potential $\Phi_0(x)$. We used simple initial potentials: either $\Phi_0(x) = \text{const}$, or a single-well potential $\Phi_0(x) = -\log(\cos^2(2\pi x/L)))$, which, respectively, correspond to $p_{\text{eq}} = \text{const}$ and $p_{eq} = \cos^2(2\pi x/L)$ ($L$ is the domain size of $x$). These two potentials minimize the model complexity (see Section *****) for the case of Neumann and Dirichlet boundary conditions, respectively. On each iteration we calculate $\mathscr{L}$ through the forward pass Eq. (13), and the variational derivative $\delta\mathscr{L}/\delta F$ using Eq. (36). Then the force is updated as:

$$F_n(x) = F_{n-1}(x) - \gamma\frac{\delta\mathscr{L}}{\delta F}, \tag{38}$$

where $\gamma$ is a learning rate [20]. In this work, the learning rate was constant over the gradient-descent iterations, but its value was different across simulations.

On each iteration, the potential is calculated from the force by integration: $\Phi_n(x) = -\int F_n(x')dx'$. An arbitrary integration constant was chosen such as:

$$\int_{-1}^{1}\exp(-\Phi_n(x))dx = 1, \tag{39}$$

which supports a convenient relationship $p_{\text{eq}} = \exp(-\Phi(x))$.

#### 1.5 Optimization of regularized likelihood

Overfitting in model selection is a general phenomenon that affects any model selection procedure over a finite data sample [26]. In particular, we verified that overfitting in model selection occurs with an explicit regularization of the likelihood. Regularization methods modify the optimization objective by adding to the likelihood a term that discourages overly complex models:

$$-\ln\mathscr{L}\left[Y(t)|\Phi(x)\right] - \eta S[\Phi(x)], \tag{40}$$

where $\eta$ is a hyperparameter.

As a regularizer, we use the trajectory entropy functional $S[\Phi(x)]$ [27]. Maximizing the trajectory entropy $S[\Phi(x)]$ discourages complex trajectories.The trajectory entropy is defined as a

Kullback-Leibler divergence between the distributions $P[X(t)]$ and $Q[X(t)]$. The former is the probability distribution of trajectories generated from the model with the potential $\Phi(x)$. The latter is the probability distribution of trajectories generated from the free diffusion (a model with a constant potential):

$$S[\Phi(x)] = -\int_0^{t_{\mathrm{obs}}} DX(t) P[X(t)] \ln \frac{P[X(t)]}{Q[X(t)]}. \tag{41}$$

The trajectory entropy can be expressed through the parameters of the Langevin dynamics [27] (here we only consider the terms that depend on the potential):

$$S[\Phi(x)] \propto -\int \left(\frac{d\Phi}{dx}\right)^2 e^{-\Phi} dx. \tag{42}$$

The hyperparameter $\eta$ controls the balance between the data-likelihood and regularizer: too small $\eta$ does not prevent overfitting, whereas too large $\eta$ results in ignoring the data and converging to the regularizer minimum. The optimal $\eta$ is usually chosen by evaluating models obtained with different $\eta$ on a validation set and selecting $\eta^*$ with the best generalization [18]. On some realizations of the training and validation data, the model at $\eta^*$ matches the ground-truth (fig. S5, *lower row*), whereas on other realizations, the model at $\eta^*$ exhibits spurious features (fig. S5, *upper row*).

For optimization with the gradient descent, we need to calculate the variational derivative of the regularizer w.r.t. the driving force $\delta S/\delta F$. Since $F(x) = p'_{\mathrm{eq}}(x)/p_{\mathrm{eq}}(x)$, Eq. (42) can be written as:

$$S[\Phi(x)] = -\int_{-1}^{1} \frac{p'^2_{\mathrm{eq}}(x)}{p_{\mathrm{eq}}(x)} dx. \tag{43}$$

Using variational chain rule [30]:

$$\frac{\delta S}{\delta F(y)} = \int_{-1}^{1} \frac{\delta S}{\delta p_{\mathrm{eq}}(s)} \frac{\delta p_{\mathrm{eq}}(s)}{\delta F(y)} ds. \tag{44}$$

The first term is calculated from Eq. (43) using the Euler-Lagrange equation:

$$\frac{\delta S}{\delta p_{\mathrm{eq}}(s)} = -\frac{p'^2_{\mathrm{eq}}(s)}{p^2_{\mathrm{eq}}(s)} + 2\frac{p''_{\mathrm{eq}}(s)}{p_{\mathrm{eq}}(s)}. \tag{45}$$

The second term is already calculated, see Eq. (33). Substituting these results, we obtain:

$$\begin{aligned} \frac{\delta S}{\delta F(y)} &= -\int_{-1}^{1} \left(\frac{p'^2_{\mathrm{eq}}(s)}{p_{\mathrm{eq}}(s)} - 2p''_{\mathrm{eq}}(s)\right) \left(H(s-y) - 1 + \int_{-1}^{y} p_{\mathrm{eq}}(s')ds'\right) ds = \\ &= -\int_{-1}^{1} \left(\frac{p'^2_{\mathrm{eq}}(s)}{p_{\mathrm{eq}}(s)} - 2p''_{\mathrm{eq}}(s)\right) \left(-H(y-s) + \int_{-1}^{y} p_{\mathrm{eq}}(s')ds'\right) ds = \\ &-\int_{-1}^{y} \left(-\frac{p'^2_{\mathrm{eq}}(s)}{p_{\mathrm{eq}}(s)} + 2p''_{\mathrm{eq}}(s) + p_{\mathrm{eq}}(s)\left[\int_{-1}^{1} \left(\frac{p'^2_{\mathrm{eq}}(s')}{p_{\mathrm{eq}}(s')} - 2p''_{\mathrm{eq}}(s')\right) ds'\right]\right) ds. \end{aligned} \tag{46}$$

#### 1.6 Model complexity

We define model complexity as the negative trajectory entropy:

$$\mathcal{M} = -S[\Phi(x)]. \tag{47}$$

The model complexity is always positive. Qualitatively, it reflects the structure of the observed trajectories: the models that produce more structured trajectories have higher complexity. Given a particular potential $\Phi(x)$, the model complexity is computed using Eqs. (42),(47).

#### 1.7 Extensions of the non-parametric framework

The non-parametric framework can be easily generalized for the simultaneous inference of $\Phi(x)$, $f(x)$, and $D$ (and even for multiple neurons with diverse firing rate functions $f_1(x)$, $f_2(x)$, ... that are coupled through the same latent dynamical system). In this case, all of these quantities are updated with the gradient-descent:

$$\begin{aligned} D_n &= D_{n-1} - \gamma_D \frac{\partial \mathscr{L}}{\partial D}, \\ f_{i,n}(x) &= f_{i,n-1}(x) - \gamma_f \frac{\delta \mathscr{L}}{\delta f_i}. \end{aligned} \tag{48}$$

The analytical expressions for the likelihood derivatives w.r.t. $D$ and $f_i(x)$ can be calculated by following exactly the same steps, as for the derivative with respect to potential, see Section 1.3. In addition, it is straight-forward to extend the framework for multiple latent dimensions. For example, in the case of two-dimensions the expressions for the likelihood and its derivatives are similar, while the dimensionality of matrices and vectors become $N^2 \times N^2$ and $N^2$ correspondingly (compared to $N \times N$ and $N$ in the case of one dimension).

#### 1.8 Simulation parameters

The full list of simulation parameters is provided in Table 1.

| parameter | value | description |
|---|---|---|
| Optimization | | |
| $\gamma$ | $5 \cdot 10^{-2}$ — $5 \cdot 10^{-4}$ | GD learning rate |
| max_it | 100000 | maximum number of iterations |
| $\eta$ | 0 — 0.1 | regularization strength |
| $\Phi_0(x)$ | $-\log\left(\cos^2 \pi x/2\right), \mathrm{const}$ | initial guess |
| $D$ | 2 — 10 | noise magnitude |
| $f(x)$ | $100(x+1)$ | firing rate function for synthetic data, Hz |
| $f(x)$ | $200(x+1)$ | firing rate function for real data, Hz |
| Spectral Elements Method | | |
| $x_{\text{begin}}$ | -1 | left boundary of the latent space |
| $x_{\text{end}}$ | 1 | right boundary of the latent space |
| $Nv$ | 64 | number of eigenfunctions of $\boldsymbol{\mathcal{H}}$ |
| bnd_cnd | Neumann | boundary conditions |
| $N_e$ | 256 | number of elements |
| $N_p$ | 8 | number of grid points in each element |
| Synthetic data generation | | |
| $T$ | 10 —1000 | trial duration, s |
| $\Delta t$ | 0.001 — 0.0001 | time bin for latent trajectory generation, s |

Table 1: **Simulation parameters**

#### 1.9 Simulations with different dynamics and data amount

We performed multiple simulations to test how the probability of overfitting in model selection depends on the data amount and on the complexity of the ground-truth dynamics. The summary

| Ground-truth model | Number of spikes in data | | |
|---|---|---|---|
| | $1,000$ | $10,000$ | $100,000$ |
| Single well | 7/3/0 | 7/3/0 | 9/1/0 |
| Double well | 6/1/3 | 6/3/1 | 10/0/0 |
| Triple well | 0/2/8 | 2/1/7 | 9/1/0 |

Table 2: **Summary of simulation outcomes.**

of simulation outcomes is presented in the Table 2. We used nine simulation settings, which used a single-, double- or triple-well ground-truth potential, each with 1,000, 10,000 and 100,000 spikes in the data sample. For each setting we performed 10 simulations on 10 independent data samples generated from the same ground-truth. Each of the data samples were split into training and validation sets of equal size. In each of the 90 independent simulations, we selected the model with the best generalization, i.e. at the minimum of the validated negative log-likelihood. We compared the model with the best generalization to the corresponding ground-truth model, and labeled the simulation outcome as consistent, overfitted or underfitted. For example, 2/3/5 in the Table 2 means that for a given simulation setting, 2 outcomes were consistent with the ground truth, 3 were overfitted and 5 were underfitted. The simulation outcome was labeled as consistent with the ground-truth if the selected model tightly overlapped with the ground-truth model. The simulation outcome was labeled as overfitted if the selected model exhibited spurious features. The simulation outcome was labeled as underfitted if some of the ground-truth features were not discovered. From the Table 2 it is clear that increasing the data amount increases the probability of consistent outcome, and thus decreases the probability of overfitting or underfitting. However, the probability of overfitting in model selection is still substantial even for ~10,000 spikes, which is similar to what is available in neurophysiological experiments. Simulations results from the cells highlighted in grey are shown in figs. S7, S8.

### 2 Materials and methods: Additional details

#### 2.1 Eigenvector-eigenvalue expansion of the Fokker-Planck operator

Consider Eq. (6) and let $\hat{\boldsymbol{\mathcal{H}}} = \hat{\boldsymbol{\mathcal{H}}}_{\mathbf{0}} + \hat{\boldsymbol{\mathcal{H}}}_{\boldsymbol{I}}$, where:

$$\hat{\boldsymbol{\mathcal{H}}}_{\mathbf{0}} = \left( D\frac{\partial}{\partial x}F(x) - D\frac{\partial^2}{\partial x^2} \right), \quad \hat{\boldsymbol{\mathcal{H}}}_{\boldsymbol{I}} = f(x). \tag{49}$$

Here $\hat{\boldsymbol{\mathcal{H}}}_{\mathbf{0}}$ is the original Fokker-Planck operator that propagates probability density $p(x,t)$ according to the Langevin dynamics, and the additional contribution $\hat{\boldsymbol{\mathcal{H}}}_{\boldsymbol{I}}$ accounts for the fact that no spikes were observed during a time period between any two adjacent spikes. We denote the equilibrium probability density of the operator $\hat{\boldsymbol{\mathcal{H}}}_{\mathbf{0}}$ as $p_{\mathrm{eq}}(x)$, and the scaled version as $\rho_{\mathrm{eq}}(x) = \sqrt{p_{\mathrm{eq}}(x)}$. This quantity is related to the potential via $p_{\mathrm{eq}}(x) = \exp(-\Phi(x))$, provided $\Phi(x)$ is normalized as in Eq. (39).

In terms of the potential, the operator $\hat{\boldsymbol{\mathcal{H}}}_{\mathbf{0}}$ can be written as:

$$\hat{\boldsymbol{\mathcal{H}}}_{\mathbf{0}} = -\frac{\partial}{\partial x}De^{-\Phi(x)}\frac{\partial}{\partial x}e^{\Phi(x)}. \tag{50}$$

To take advantage of the useful properties of Hermitian operators, we multiply Eq. (6) by

$\exp(\Phi(x)/2)$:

$$e^{\Phi(x)/2}\frac{\partial p}{\partial t} = -\underbrace{e^{\Phi(x)/2}\hat{\boldsymbol{\mathcal{H}}}e^{-\Phi(x)/2}}_{\mathcal{H}}\underbrace{e^{\Phi(x)/2}p}_{\rho}. \tag{51}$$

Here we introduced a new Hermitian operator $\boldsymbol{\mathcal{H}} = \exp(\Phi(x)/2)\hat{\boldsymbol{\mathcal{H}}}\exp(-\Phi(x)/2)$. This operator acts on $\rho(x,t) = p(x,t)/\rho_{eq}(x)$, where $\rho_{eq}(x) = \exp(-\Phi(x)/2) = \sqrt{p_{eq}(x)}$. Similar to the original operator, we decompose $\boldsymbol{\mathcal{H}} = \boldsymbol{\mathcal{H}_0} + \boldsymbol{\mathcal{H}_I}$, where

$$\boldsymbol{\mathcal{H}_0} = e^{\Phi(x)/2}\hat{\boldsymbol{\mathcal{H}}}_{\mathbf{0}}e^{-\Phi(x)/2}, \qquad \hat{\boldsymbol{\mathcal{H}}}_{\boldsymbol{I}} = \boldsymbol{\mathcal{H}_I}. \tag{52}$$

Following [15], we first solve the eigenvalue-eigenvector problem for the operator $\boldsymbol{\mathcal{H}_0}$:

$$\boldsymbol{\mathcal{H}_0}\Psi_0(x) = -e^{\Phi(x)/2}\frac{\partial}{\partial x}De^{-\Phi(x)}\frac{\partial}{\partial x}e^{\Phi(x)/2}\Psi_0(x) = \lambda_0\Psi_0(x). \tag{53}$$

Multiplying both sides by $\exp(-\Phi(x)/2)$ from the left and letting $\phi_0(x) = \Psi_0(x)\exp(\Phi(x)/2) = \Psi_0(x)/\sqrt{p_{eq}(x)}$, we obtain:

$$-\frac{\partial}{\partial x}De^{-\Phi(x)}\frac{\partial}{\partial x}\phi_0(x) = \lambda_0 p_{eq}(x)\phi_0(x). \tag{54}$$

This problem is discretized in the SEM basis, see below. Since Eq. (54) is of the second order, one has to impose two boundary conditions. Assuming reflective boundaries for the original probability distribution $p(x,t)$ (zero probability flux), it can be shown [31] that our problem satisfies backward Fokker-Planck equation and obeys Neumann (zero-derivative) boundary conditions. Eigenvalues $\boldsymbol{\lambda_0}$ and eigenvectors $\phi_{0,i}(x)$ of the discretized problem are obtained with Python solver *scipy.linalg.eigh*. This basis is truncated to the first 64 basis functions. After that the eigenvectors of $\boldsymbol{\mathcal{H}_0}$ are obtained by $\Psi_0(x) = \phi_0(x)\rho_{eq}(x)$. Next the eigenvalue problem for the full operator $\boldsymbol{\mathcal{H}}$ is solved in the $\Psi_0$ basis. Similar to [15], the matrix form of this problem is $\boldsymbol{K}\hat{\Psi} = \lambda\hat{\Psi}$, where:

$$K_{ij} = \langle\Psi_{0,i}|\boldsymbol{\mathcal{H}_0} + \boldsymbol{\mathcal{H}_I}|\Psi_{0,j}\rangle = \lambda_{0,i}\delta_{ij} + \langle\Psi_{0,i}|f(x)|\Psi_{0,j}\rangle = \lambda_{0,i}\boldsymbol{I} + \boldsymbol{Q}_0^T\boldsymbol{W}f(x)\boldsymbol{Q}_0. \tag{55}$$

Eigenvectors of the operator $\boldsymbol{\mathcal{H}}$ in the original SEM basis are then obtained by the rules of basis transformation: $\boldsymbol{Q} = \boldsymbol{Q}_0\hat{\boldsymbol{Q}}$, where $\boldsymbol{Q}_0$ and $\hat{\boldsymbol{Q}}$ are the eigenvector matrices of $\boldsymbol{\mathcal{H}_0}$ and $\boldsymbol{K}$, correspondingly.

#### 2.2 Spectral elements method

Following [15], we use the Spectral Elements Method (SEM) for discretization of the eigenvalue problem Eq. (54). This method is numerically efficient due to sparse discretization and has exponential precision as other spectral methods. SEM is briefly outlined here, while the details can be found in [32].

Consider the following Sturm-Liouville problem for the function $u(x)$:

$$\begin{gathered}-\frac{d}{dx}\left(p(x)\frac{du(x)}{dx}\right) + q(x)u(x) = \lambda w(x)u(x), \qquad a < x < b,\\ A_1u(a) + A_2u'(a) = 0, \quad B_1u(b) + B_2u'(b) = 0.\end{gathered} \tag{56}$$

Using the weak formulation, we multiply both sides of Eq. (56) by a test function $\Xi_i(x)$ and integrate:

$$\int_a^b\left(-\frac{d}{dx}\left(p(x)\frac{du(x)}{dx}\right)\right)\Xi_i(x)dx + \int_a^b q(x)u(x)\Xi_i(x)dx = \int_a^b \lambda w(x)u(x)\Xi_i(x)dx. \tag{57}$$

The first term can be integrated by parts:

$$\left[-p(x)\frac{du(x)}{dx}\Xi_i(x)\right]_a^b + \int_a^b p(x)\frac{du(x)}{dx}\frac{d\Xi_i(x)}{dx}dx + \int_a^b q(x)u(x)\Xi_i(x)dx = \int_a^b \lambda w(x)u(x)\Xi_i(x)dx. \tag{58}$$

Next, we assume a finite number of collocation points $x_i$, $i = 1, 2, ..., N$ and use (yet unspecified) basis that consists of $N$ basis functions $\Psi_i(x)$. Expanding $u(x)$ with this basis, we get:

$$u(x) = \sum_{i=1}^{N} u_i \Psi_i(x), \qquad \frac{du(x)}{dx} = \sum_{i=1}^{N} u_i \frac{d\Psi_i(x)}{dx}. \tag{59}$$

By using the basis function instead of test functions and substituting the above expansions into Eq. (58), we get:

$$\begin{aligned}\left[-p(x)\frac{du(x)}{dx}\Psi_i(x)\right]_a^b + \sum_j \int_a^b p(x)u_j\frac{d\Psi_j(x)}{dx}\frac{d\Psi_i(x)}{dx}dx + \sum_j \int_a^b q(x)u_j\Psi_j(x)\Psi_i(x)dx = \\ = \sum_j \int_a^b \lambda w(x)u_j\Psi_j(x)\Psi_i(x)dx.\end{aligned} \tag{60}$$

Using our collocation points and replacing integrals with sums (and weights $\rho_k$), we obtain:

$$\begin{aligned}\left[-p(x)\frac{du(x)}{dx}\Psi_i(x)\right]_a^b + \sum_j\sum_k \rho_k p(x_k)u_j\frac{d\Psi_j(x_k)}{dx}\frac{d\Psi_i(x_k)}{dx} + \sum_j\sum_k \rho_k q(x_k)u_j\Psi_j(x_k)\Psi_i(x_k) = \\ = \lambda\sum_j\sum_k \rho_k w(x_k)u_j\Psi_j(x_k)\Psi_i(x_k).\end{aligned} \tag{61}$$

At this point we need to specify the basis functions and collocation points. Following [32], we use Gauss-Lobatto-Legendre (GLL) grid and Lagrange interpolation polynomials as basis functions. The GLL grid specifies the collocation points $x_k$ and integration weights $\rho_k$:

$$\begin{aligned} x_k: &\quad x_1 = -1, \quad x_N = 1, \text{ and all zeros of } L'_{N-1}(x), \\ \rho_k &= \frac{2}{N(N-1)}\frac{1}{[L_{N-1}(x_k)]^2}. \end{aligned} \tag{62}$$

Lagrange interpolation polynomials are defined as:

$$\Psi_j(x) = \prod_{\substack{1<m<N \\ m\neq j}} \frac{x - x_m}{x_j - x_m}. \tag{63}$$

They have a useful property $\Psi_j(x_i) = \delta_{ij}$, hence the expansion coefficients in Eq. (59) are trivial: $u_i = \Psi_i(x_i)$. By introducing a differentiation matrix $D_{ij} = d\Psi_j(x_i)/dx$, Eq. (61) can be written as:

$$\left[-p(x)\frac{du(x)}{dx}\Psi_i(x)\right]_a^b + \sum_j\left(\sum_k \rho_k p(x_k)D_{kj}D_{ki}\right)u(x_j) + \sum_j\left(\rho_i q(x_i)\delta_{ij}\right)u(x_j) = \lambda\sum_j\left(\rho_i w(x_i)\delta_{ij}\right)u(x_j). \tag{64}$$

This can be written in matrix-vector form:

$$\left[-p(x)\frac{du(x)}{dx}\Psi_i(x)\right]_a^b + \mathbf{H}\tilde{\mathbf{u}} = \lambda\mathbf{M}\tilde{\mathbf{u}}. \tag{65}$$

Here the stiffness $\boldsymbol{H}$ and mass $\boldsymbol{M}$ matrices are defined as:

$$H_{ij} = \sum_k \left(D^T_{ik}\rho_k p(x_k) D_{kj}\right) + \rho_i q(x_i)\delta_{ij}, \qquad M_{ij} = \rho_i w(x_i)\delta_{ij}. \tag{66}$$

For the chosen grid and basis functions, the differentiation matrix is known [32]:

$$D_{ij} = \frac{d\Psi_j(x_i)}{dx} = \begin{cases} \frac{L_{N-1}(x_i)}{L_{N-1}(x_j)}\frac{1}{x_i - x_j}, & i \neq j, \\ -\frac{N(N-1)}{4}, & i = j = 1, \\ \frac{N(N-1)}{4}, & i = j = N, \\ 0, & \text{otherwise.} \end{cases} \tag{67}$$

By comparing Eq. (56) with our problem Eq. (54), we see that in Sturm-Liouville notations:

$$p(x) = De^{-\Phi(x)} = Dp_{\mathrm{eq}}(x), \qquad q(x) = 0, \qquad w(x) = p_{\mathrm{eq}}(x). \tag{68}$$

Our goal is to discretize Eq. (65) with Neumann boundary conditions. These conditions imply that the boundary term in Eq. (65) is zero.To solve Eq. (65), we split our domain into $N_e$ equal elements, each of which contains $N_p$ points, including the boundary points, such that each of the adjacent elements have only one point in common. Next, a single element is mapped into an interval $[-1;1]$ using affine mapping. After that the local stiffness and mass matrices are calculated on a single element using Eq. (66). To construct the global $\boldsymbol{H}$ and $\boldsymbol{M}$ for the entire domain, we map the local matrices back to the $x$-domain and patch the pieces together, see [32] for details. The resulting matrices have block-diagonal structure, where the width of the block is equal to the number of collocation points on a single element, $N_p$. After that, the discretized eigenvector-eigenvalue problem was solved with *scipy.linalg.eigh* python solver. For the values of the parameters refer to Table S1.

#### 2.3 Antiderivatives using Lagrange polynomials

In this work we used Lagrange polynomials basis, $f(x) = \sum_i f(x_i)\Psi_i(x)$, where $\Psi_i(x)$ is given by (63). To calculate the antiderivative of any function $F(x) = \int_{-1}^{x} f(x)dx$, we use our Lagrange expansion of $f(x)$ and integrate each of the Lagrange polynomials analytically. The integration matrix is thus:

$$B_{lk} = \int_{-1}^{x_l} \Psi_k(x')dx'.$$

With this notation, the antiderivative is calculated with a matrix-vector multiplication:

$$F(x_l) = \int_{-1}^{x_l} f(x')dx' = \sum_{k=1}^{N} f(x_k)\int_{-1}^{x_l}\Psi_k(x')dx' = (\boldsymbol{Bf})_l\,.$$

The integration matrix can be calculated from definition of Lagrange polynomials (63):

$$\boldsymbol{B}_{lk} = \frac{1}{D_k}\int_{-1}^{x_l}\prod_{i\neq k}(x' - x_i)dx', \qquad D_k = \prod_{i\neq k}(x_k - x_i).$$

Expanding the parentheses under the integral, we obtain a polynomial of degree $(N-1)$, so that the result will be a polynomial of degree $N$. The coefficients near each of $x^k$ will be the sum of all possible $k$-wise products. After some algebra the final result is given by:

$$\boldsymbol{B}_{lk} = \frac{1}{D_k}\sum_{n=1}^{N} C_{nk}x_l^n + C_{0k}, \qquad C_{nk} = \frac{(-1)^{\mathrm{mod(N-n,2)}}}{n}\mathrm{Com(N-n, \{X \setminus x_k\})}. \tag{69}$$

Here mod is the reminder after division, $\{X \setminus x_k\}$ is a set of all the grid points except for $x_k$, and $\mathrm{Com(n, \{X \setminus x_k\})}$ denotes the sum of all possible $n$-wise products of a given set, for example:

$$\begin{aligned}
&Com(0, \{x_1, x_2, x_3, x_4\}) = 1 \\
&Com(1, \{x_1, x_2, x_3, x_4\}) = x_1 + x_2 + x_3 + x_4 \\
&Com(2, \{x_1, x_2, x_3, x_4\}) = x_1x_2 + x_1x_3 + x_1x_4 + x_2x_3 + x_2x_4 + x_3x_4 \\
&Com(3, \{x_1, x_2, x_3, x_4\}) = x_1x_2x_3 + x_1x_2x_4 + x_1x_3x_4 + x_2x_3x_4 \\
&Com(4, \{x_1, x_2, x_3, x_4\}) = x_1x_2x_3x_4.
\end{aligned} \tag{70}$$

Finally, $C_{0k}$ is an arbitrary integration constant. We set $F(-1) = 0$, which can be implemented by setting the first row of integration matrix to zero.

To adapt this method for finite element grid, we calculated the integration matrix on a single element, scaled it back on a full domain and patched the global integration matrix by stacking the local ones on the main diagonal. To account for an additive property of integration (accumulation of the integration results when moving from the left to the right), we added a bias vector $\boldsymbol{u}$. This vector accumulates the result of integration from the previous elements (that are located to the left of the current element). The final result is:

$$\begin{aligned}
&\tilde{F} = \boldsymbol{Bf}, \\
&u = \{\underbrace{0, 0, ..., 0}_{N_p}, \underbrace{\tilde{F}_{N_p-1}, \tilde{F}_{N_p-1}, ...\tilde{F}_{N_p-1}}_{N_p-1}, \underbrace{\tilde{F}_{N_p-1} + \tilde{F}_{2(N_p-1)}, ..., \tilde{F}_{N_p-1} + \tilde{F}_{2(N_p-1)}}_{N_p-1}, ...\}, \\
&F = \boldsymbol{Bf} + \boldsymbol{u}.
\end{aligned} \tag{71}$$

Alternatively, the bias can be absorbed into the differentiation matrix, which would make the differentiation matrix lower triangular instead of sparse.

### 3 Supplementary figures

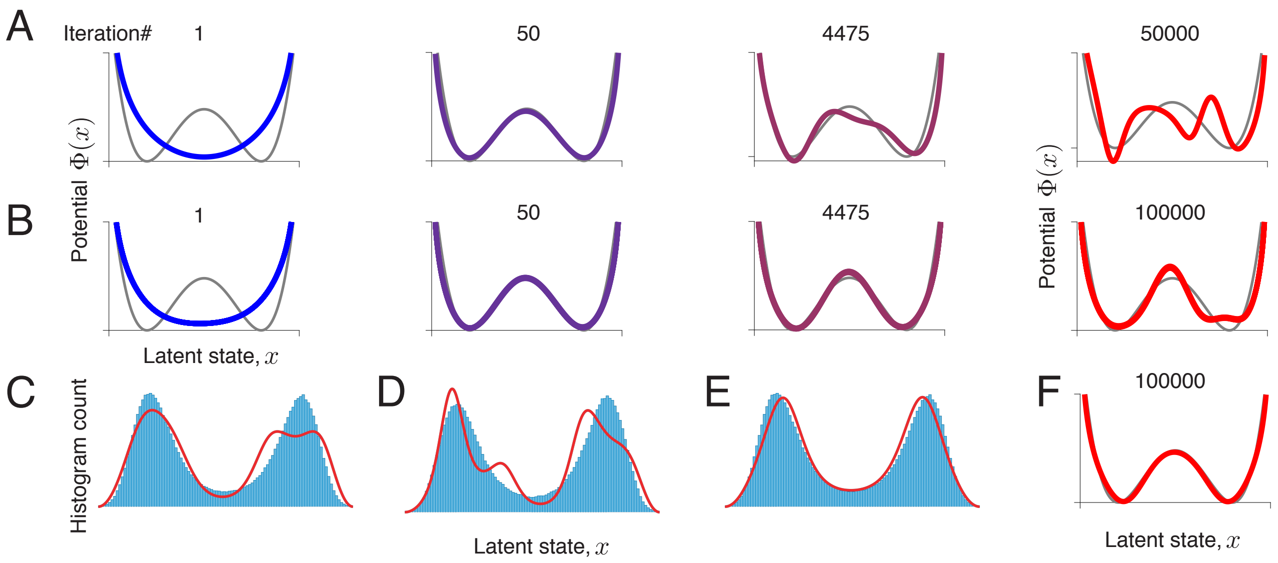

Figure 1: **Overfitting does not occur with infinite data.** (**A**) A series of models produced by gradient-descent, when the same finite set of data is used throughout the optimization. Substantial overfitting is observed. (**B**) Same as A, but optimization is performed with spikes resampled on each iteration of gradient-descent from a fixed latent trajectory (latent trajectory is the same as in A). Overfitting is still observed. (**C**) Histogram of the latent trajectory (normalized as probability density) and discovered equilibrium probability density (at iteration 100,000) from the simulation in B. Overfitted model contains features that are not presented in the latent trajectory. (**D**) Same as C, but for a different latent trajectory. Spurious features in C and D are different. (**E**) Same as C, but with resampling both the latent trajectory and spikes on each iteration of gradient-descent. No signs of overfitting are observed. (**F**) After 100,000 iterations, the inferred potential for the simulation in E still perfectly matches the ground truth. These simulations confirm that overfitting emerges largely due to Poisson noise (compare A and B), although noise in the latent trajectory also contributes to overfitting (compare B and F). No overfitting occurs when both spikes and latent trajectory are resampled on each gradient descent iteration. In all panels, the training and validation datasets (with the size of ∼10,000 spikes) were generated from a double-well potential.

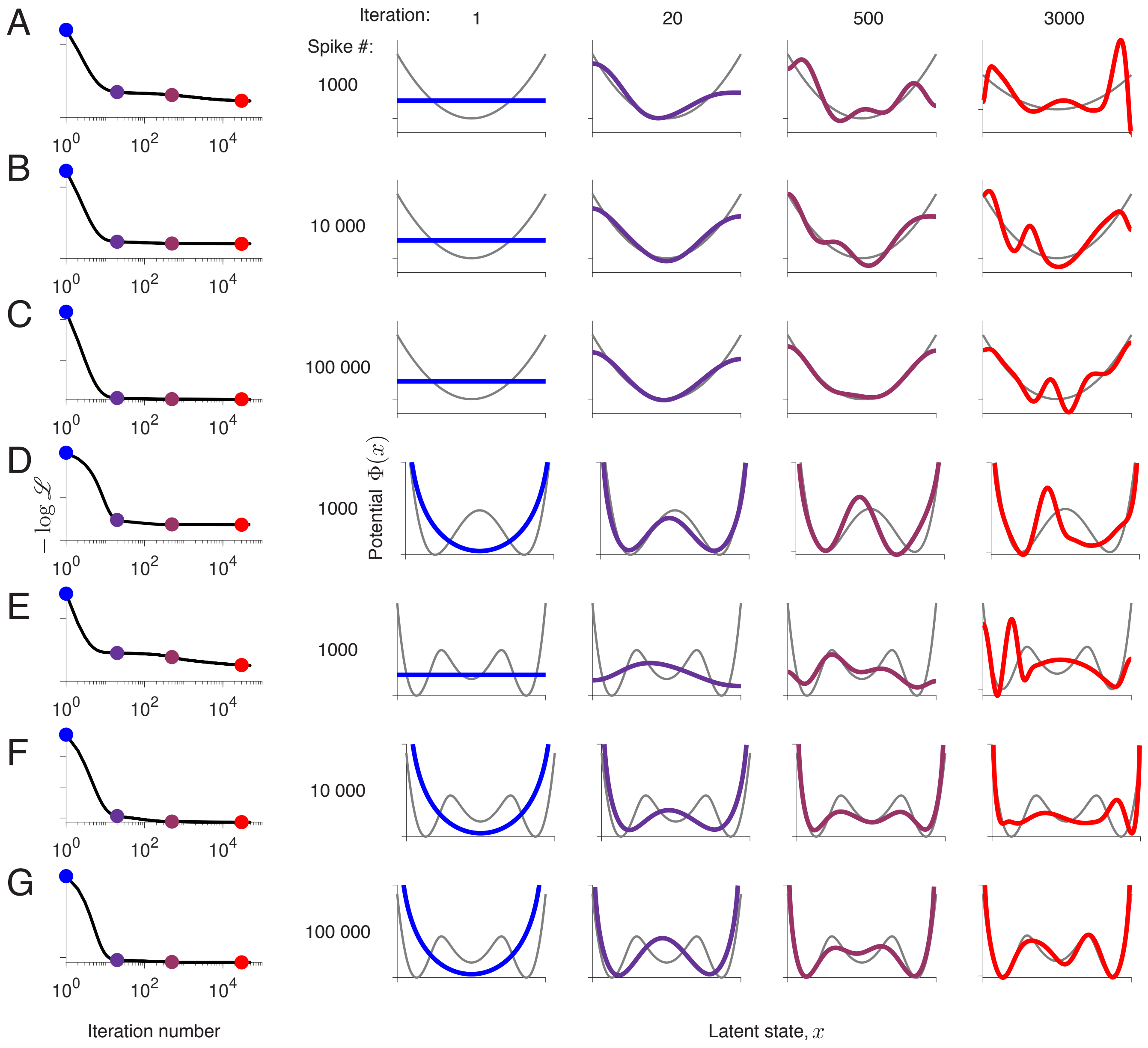

Figure 2: **Overfitting is universally observed for different dynamics, data amount and hyperparameters.** Negative log-likelihood $-\log \mathscr{L}$ over iterations of gradient descent (left), and fitted potentials on selected iterations (right, colors correspond to the dots on the likelihood plot). The rows show simulations with different ground-truth dynamics, size of the data sample and initial condition for optimization ($\Phi_0(x)$). (**A**) Single-well potential, 1,000 spikes, and $\Phi_0(x)$ = const. (**B**) Single-well potential, 10,000 spikes, and $\Phi_0(x)$ = const. (**C**) Single-well potential, 100,000 spikes, and $\Phi_0(x)$ = const. (**D**) Double-well potential, 1,000 spikes, and $\Phi_0(x) = -\log(\cos^2(\pi x/2))$. (**E**) Triple-well potential, 1,000 spikes, and $\Phi_0(x)$ = const. (**F**) Triple-well potential, 10,000 spikes, and $\Phi_0(x) = -\log(\cos^2(\pi x/2))$. (**G**) Triple-well potential, 100,000 spikes, and $\Phi_0(x) = -\log(\cos^2(\pi x/2))$.

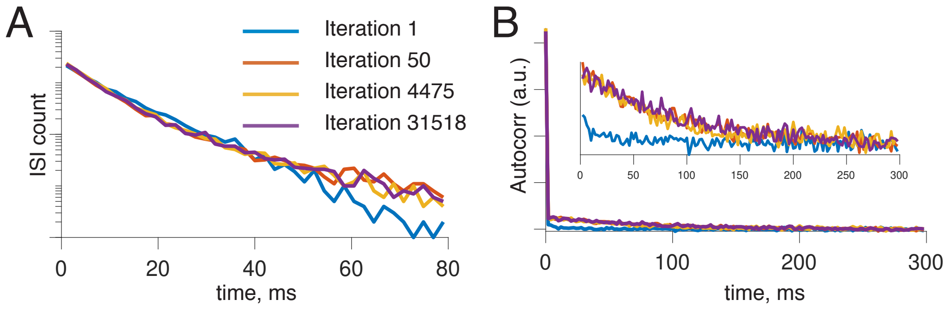

Figure 3: **Statistics of spikes in overfitted models.** Statistics of spikes are computed for the simulation in Fig. 2B,C in the main text. Synthetic spike data was generated from the fitted potentials on four selected iterations (1, 50, 4475 and 31518). Interspike interval (ISI) distributions and autocorrelation functions were computed for these data. (**A**) ISIs distributions. (**B**) Autocorrelation functions (inset: zoom on vertical axis). The spike statistics are very similar for the fitted potentials obtained on iterations 50, 4475 and 31518, despite these potentials exhibit different features (see Fig. 2B,C).

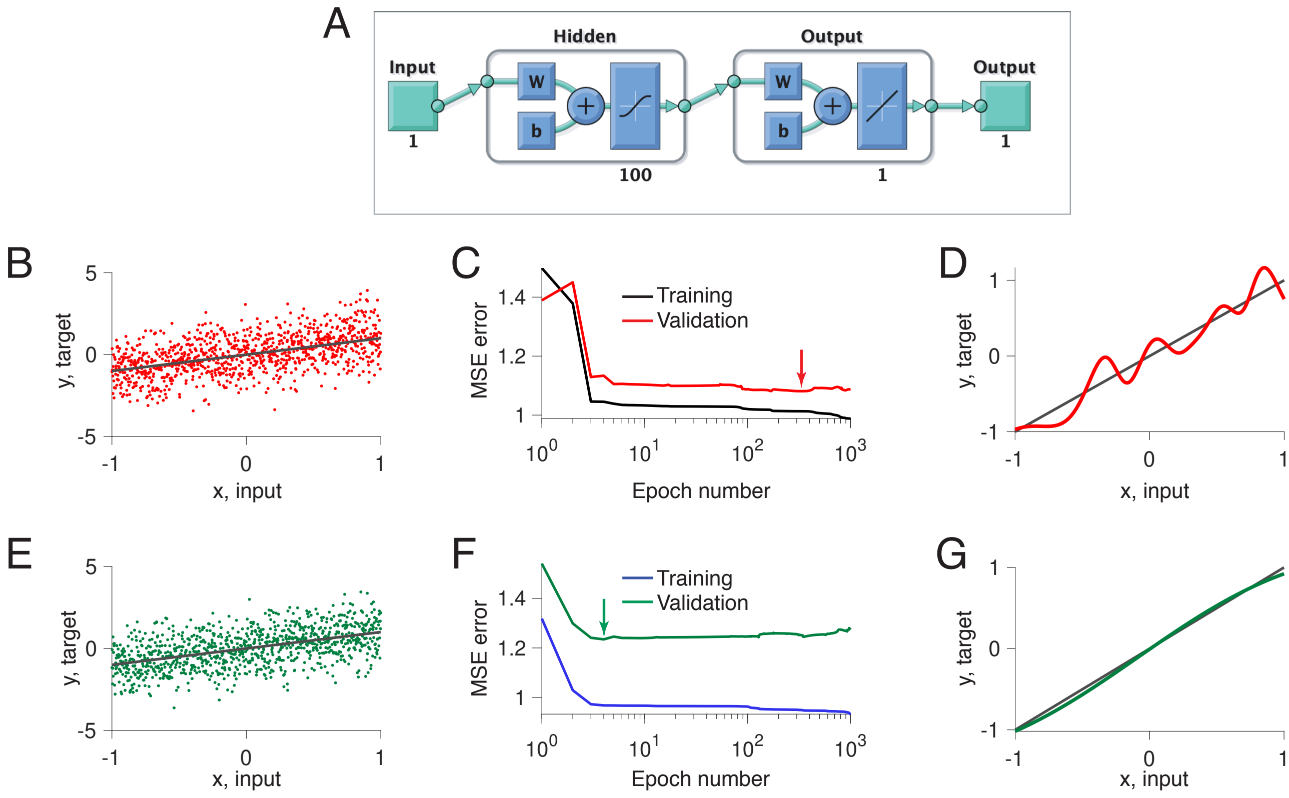

Figure 4: **A feedforward neural network exhibits generalization plateaus and overfitting in model selection.** We train a shallow feedforward neural network (1 hidden layer, 100 neurons, the total number of parameters is 301) on a regression problem. The noisy dataset of 1,000 samples was generated from a linear model $y = x + 0.2\xi$, where $\xi \sim \mathcal{N}(0, 1)$. The network was trained using Matlab Deep Learning toolbox, which runs stochastic gradient-descent with early stopping regularization. We initialized all parameters (weights and biases) from the normal distribution with zero mean and variance 0.01. We verified that our results do not dependent on a particular realization of the initial parameters. (**A**) Network architecture. (**B**) Example dataset (dots) along with the linear ground-truth model (line). (**C**) Training and validated mean squared errors (MSEs) over the optimization epochs. Long plateau in the validated MSE indicate that many models generalize equally well. Arrow indicates the minimum of the validated MSE, i.e. the model with the best generalization. (**D**) The model with the best generalization (red line, corresponds to arrow in C) contains spurious features not present in the linear ground-truth model (grey line). (**E**) Same as B but for a different data sample from the same ground-truth model. (**F**) Same as C, but for the data sample in E. (**G**) Same as D, but for the data sample in E. The model with the best generalization (green line, corresponds to arrow in F) closely matches the ground-truth model (grey line).

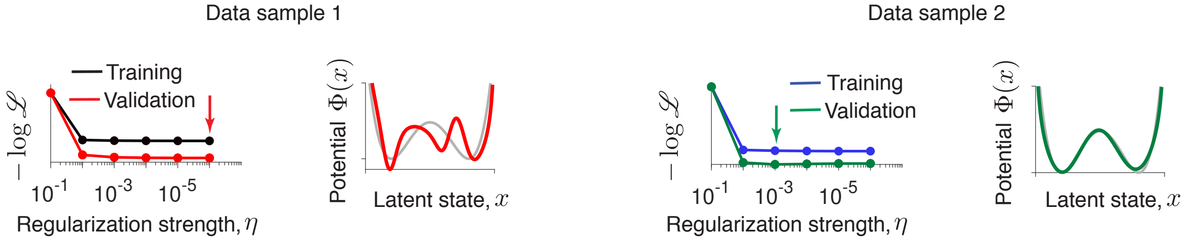

Figure 5: **Regularization does not prevent overfitting in model selection.** Optimization is performed with a regularized likelihood (see Section 1.5). Training and validated negative log-likelihood (the first and the third panels) for two data samples generated from the same ground-truth model are shown. The likelihoods are shown for the models fitted with different levels of regularization $\eta$ (x-axis). Arrows indicate the minimum of the validated negative log-likelihood, corresponding to the model with the best generalization at the optimal regularization level $\eta^*$. The corresponding potential $\Phi^*(x)$ (the second and the fourth panels, colored lines) matches the ground truth (grey) on data sample 2 and is overfitted on data sample 1.

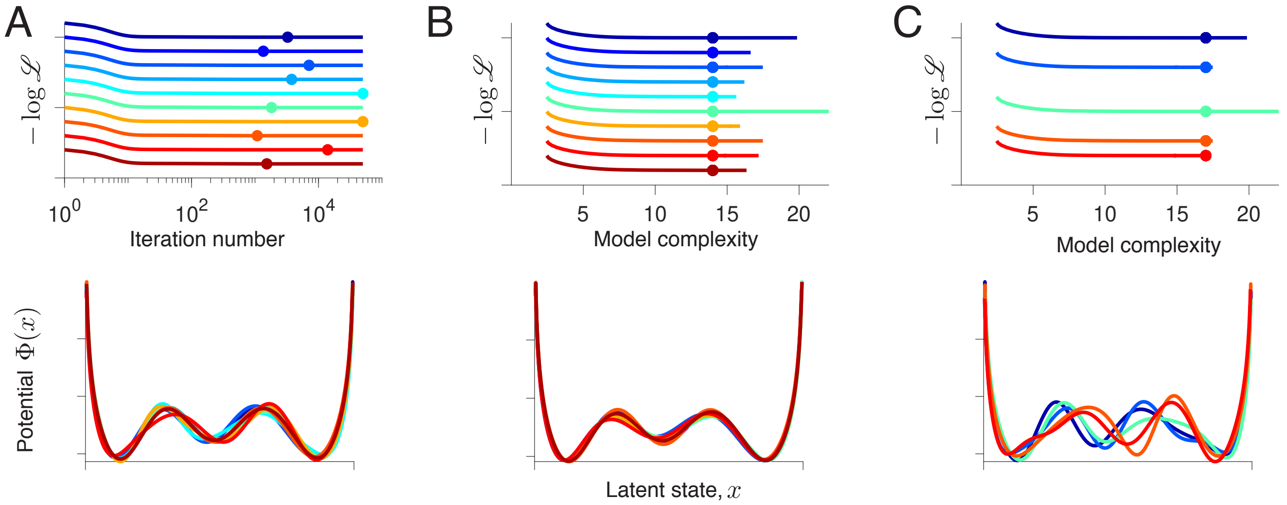

Figure 6: **Selecting the model with correct interpretation (large data amount).** Each colored line represents one out of 10 simulations, which were performed with the same settings on 10 independent data samples (triple-well ground-truth potential, 100,000 spikes). (**A**) Upper panel: Validated negative log-likelihood achieves minimum (dots) at different gradient-descent iterations on different data samples. Lowe panel: Fitted potentials selected at the minimum of the negative validated log-likelihood are largely consistent across data samples due to a very large data amount. (**B**) Upper panel: Same data as in A, but the validated negative log-likehood is plotted as a function of the model complexity. Model complexity threshold $\mathcal{M}^*$ (dots) is chosen by tracking the consistency of features along the model complexity axis. Lowe panel: The fitted potentials obtained at $\mathcal{M}^*$ are in great agreement with the ground truth. (**C**) Upper panel: Same data as in B, but a higher model complexity is chosen (dots). Lower row: At this higher model complexity, the fitted potentials are inconsistent across data samples.

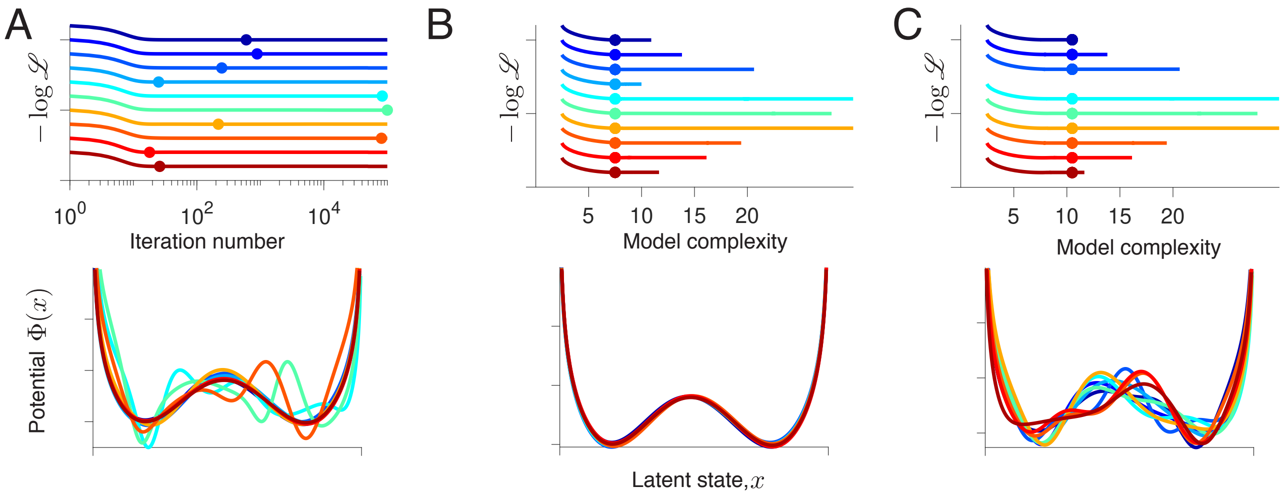

Figure 7: **Selecting the model with correct interpretation (moderate data amount).** Each colored line represents one out of 10 simulations, which were performed with the same settings on 10 independent data samples (double-well ground-truth potential, 10,000 spikes). (**A**) Upper panel: Validated negative log-likelihood achieves minimum (dots) at different gradient-descent iterations on different data samples. Lowe panel: Fitted potentials selected at the minimum of the negative validated log-likelihood are inconsistent across data samples, and many of them exhibit spurious features. (**B**) Upper panel: Same data as in A, but the validated negative log-likehood is plotted as a function of the model complexity. Model complexity threshold $\mathcal{M}^*$ (dots) is chosen by tracking the consistency of features along the model complexity axis. Lowe panel: The fitted potentials obtained at $\mathcal{M}^*$ are in great agreement with the ground truth. (**C**) Upper panel: Same data as in B, but a higher model complexity is chosen (dots). Lower row: At this higher model complexity, the fitted potentials are inconsistent across data samples.

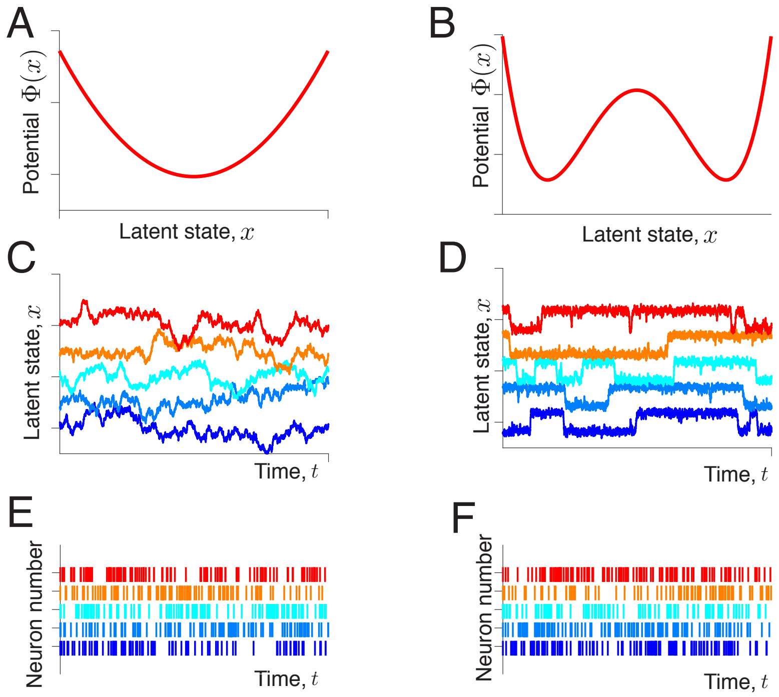

Figure 8: **Spike trains generated from the double-well and single-well models appear similar.** (**A**) Single-well potential. (**B**) Double-well potential. The potentials are chosen so that the second order spike-statistics are similar to that of the single-well potential in A. (**C**) Five independent realizations of the latent trajectories generated from a single-well potential in A. (**D**) Five independent realizations of the latent trajectories generated from a double-well potential in B. (**E**) Five spike trains generated from the latent trajectories in C. (**F**) Five spike trains generated from the latent trajectories in D.

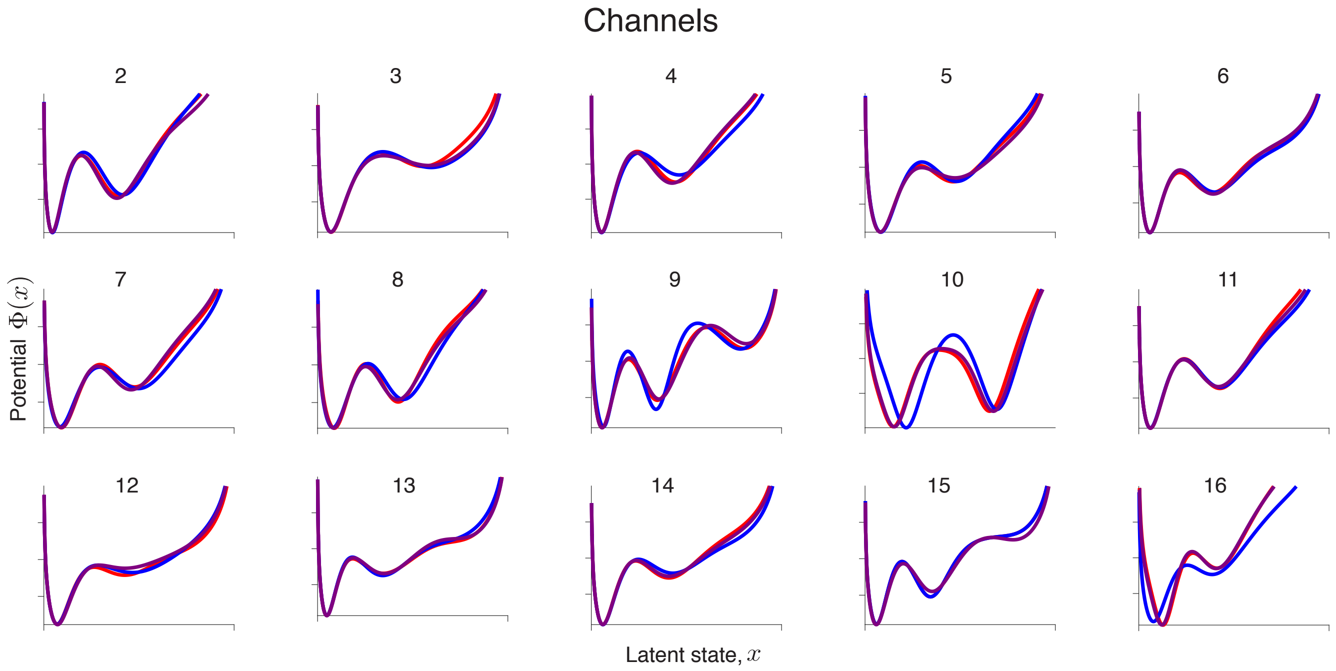

Figure 9: **Models discovered from neurophysiological spike recordings.** Each channel 1-16 is analyzed independently. For each channel we split the full data $\mathscr{D}$ in two halves $\mathscr{D}1$ and $\mathscr{D}2$, and performed optimization on three data samples: $\mathscr{D}1$, $\mathscr{D}2$ and $\mathscr{D}$. The model complexity threshold $\mathcal{M}^*$ is the maximal model complexity for which potentials are consistent across data samples $\mathscr{D}1$, $\mathscr{D}2$ and $\mathscr{D}$. Fitted potentials selected at $\mathcal{M}^*$ are shown for channels 2-16 (channel 1 is shown in Fig. 4 of the main text). The inferred double-well potential shape is highly consistent across channels.
